# Task-similarity dependent reconfiguration of compositional modules and geometry in frontal cortex

**DOI:** 10.1101/2025.11.06.687080

**Authors:** Xia Chen, Xiao Yao, Xinxin Yin, Zengcai V. Guo

## Abstract

Task rules critically shape how neural populations encode information and generalize across cognitive demands, yet the mechanisms linking rule structure, neural dynamics, and behavior remain poorly understood. Here we investigate how rule congruence modulates neural-ensemble compositionality, geometry and cross-task generalizability in mouse anterior lateral motor cortex (ALM). Mice switched between tactile and auditory delayed-response tasks where sensory-motor contingencies were either aligned (congruent) or conflicting (incongruent). Recording from 11,000 ALM neurons revealed rule dependent compositional coding of stimulus, choice, and outcome. In contrast to congruent switching, incongruent rules reduced cross-task generalizability of sensory, choice and outcome representations, expanded population dimensionality, increased reliance on nonlinear mixed-selectivity, and produced more complex coding geometry. These neural changes paralleled behavioral costs during task transitions. Moreover, congruence dependent reconfiguration of sensory-choice subspaces in ALM predicted differences in transition costs, implicating ALM in adaptively resolving conflicting contingencies. Together, these results show that task congruence regulates neural coding geometry and population dynamics, linking rule-switching complexity to dimensionality of neural resource allocation and to cognitive flexibility.

## Introduction

In a complex and changing environment, animals need to flexibly perform a wide variety of tasks. The ability to adapt to dynamic environment hinges on the brain’s capacity to reorganize neural computations across shifting task demands. The frontal cortex, a hub area for cognitive control, plays a central role in this process. Neurons in frontal cortex exhibit diverse selectivity profiles, encoding task variables including sensory cue, choice, action, and outcome often in a combinatorial way^1–8^. This property enables high-dimensional neural representations that support complex, context-dependent behaviors^5,9–11^. While neurons with nonlinear mixed selectivity (NMS) have been proposed to enhance computational capacity and flexibility^10–14^, how neural populations dynamically reconfigure across tasks, particularly under variable rule structures, remains poorly understood.

Recent network models and experiments suggest that frontal population codes are structured rather than randomly mixed. Neural network models trained on multiple cognitive tasks develop compositional architectures, where motifs for elementary computations are flexibly combined to meet task demands^15–17^. Concordantly, physiological studies indicate that representations of sensory, choice, action and outcome information are organized into functional modules embedded within low-dimensional space^18,19^. In tasks involving multiple sensory modalities, cross-modal representations with varying degrees of overlap have been observed in frontal and parietal cortices^20–28^. While shared neural representations across tasks may arise from common task components, dissimilar representations can be driven by differences in task sensory, cognitive or motor components as well as differences in task rules. Disentangling these contributions is challenging, particularly because single-neuron responses are heterogeneous and neural representations often rely on nonlinear mixtures of variables. As a result, it remains unclear how rule structure shapes the compositionality of neural ensembles, the dimensionality and geometry of neural representations, and the extent to which representations generalize across tasks.

Here, we address this gap by developing multitask behavioral paradigms in which mice performed tactile and auditory delayed-response tasks and switched between them within a session in the same behavioral apparatus. The two tasks shared temporal structure and motor planning demands, requiring the association of one of two stimuli with a directional lick after a temporal delay. Critically, we manipulated the sensory-motor mapping across two contexts: in one, tactile and auditory contingencies were congruent (aligned), while in the other they were incongruent (conflicting). Comparing effects of sensorimotor mapping across different contexts isolates the influence of rule congruence while holding constant task timing and overall sensorimotor demands.

We focus on the anterior lateral motor cortex (ALM), a frontal region critical for decision-making, motor planning, and execution^29–34^, and has recently been implicated in supporting context-dependent flexible behaviors^35–37^. Combining population-level dimensionality reduction, decoding, and cross-task generalization analyses, we asked three questions: 1) How do ALM ensembles compositionally encode stimulus, choice, and outcome across task rules? 2) How does rule congruence modulate the dimensionality and geometry of ALM population activity? 3) To what extent do these neural changes predict behavioral flexibility during task transition?

We recorded 11,000 neurons from ALM and find that rule structure exerts a systematic influence on ALM coding. Under congruent rules, ALM representations of stimulus, choice, and outcome exhibit higher cross-task generalizability and occupy lower-dimensional subspaces, consistent with efficient reuse of compositional coding primitives. In contrast, incongruent rules reduce cross-task generalizability, expand neural population dimensionality, and increase reliance on neurons with nonlinear mixed selectivity, manifesting signatures of elevated computational demands. These differences parallel behavioral costs during transitions. The degree of rule-driven reconfiguration in ALM sensory-choice subspaces predicts tasks differences in switching difficulty, highlighting ALM’s role in resolving conflicting contingencies. Together, these results advance understanding of how rule-switching complexity regulates the dimensionality of neural resource allocation and cognitive flexibility in frontal cortex.

## Results

### Congruent and incongruent rules differentially modulate task transitions

To dissect how rule-congruence shapes cross-task adaptability, we developed a sensory-modality based task-switching paradigm (Fig. 1a; Methods). Head-fixed mice performed tactile and auditory delayed-response tasks^29,38^ that require them to discriminate stimuli (whisker vibration amplitude or sound intensity) during a sample epoch, withhold licks during a delay epoch, and report decisions by licking left/right water spouts in the response epoch. Critically, mice were required to switch between tasks without explicit cues, allowing us to probe how intrinsic task structure, i.e. the sensory-motor contingency alignment, governs behavioral flexibility and neural coding. Two important features further facilitate the investigation of neural ensemble compositionality. First, the tactile and auditory tasks enable symmetric motor responses with equivalent rewards for left and right choices, allowing neural coding differentiation of choice and outcome. Second, the delay epoch separates sensory inputs from motor actions, minimizing licking-related confounds on choice-related motor planning activity.

**Fig. 1.**
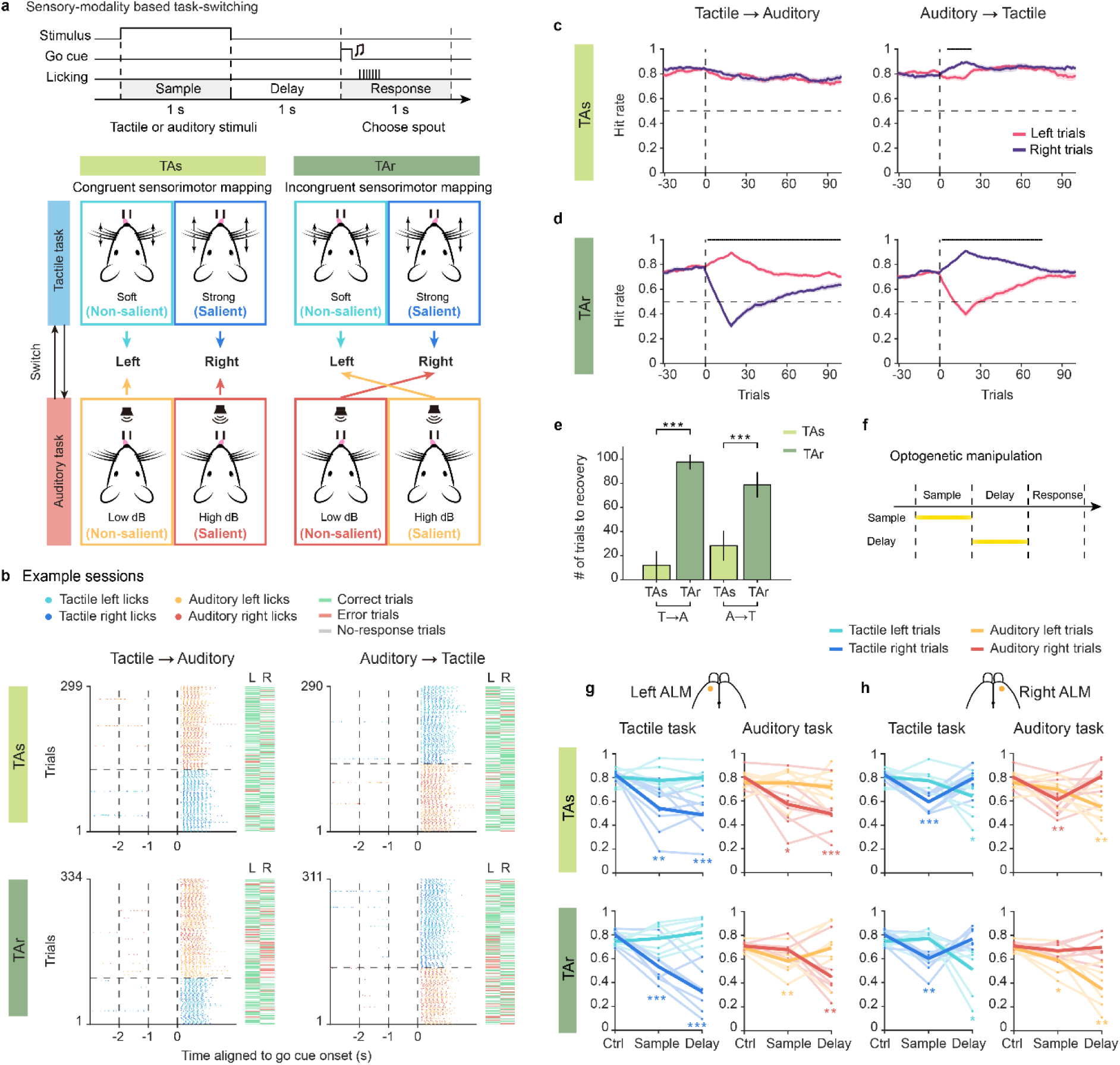
Behavioral paradigms and performance. **a**, Schematic of sensory-modality based task-switching paradigm. Mice discriminate stimulus saliency during the sample epoch and report by licking left or right spout after a delay. The tactile and auditory modality switched with blocks and the sensorimotor mapping can be congruent or incongruent. Under the tactile-auditory congruent condition (TAs), the sensorimotor mapping is similar; both strong whisker vibration and high decibel tones (salient stimuli) indicated rightward licking. Under the tactile-auditory incongruent condition (TAr), sensorimotor mapping is reversed after switching. Light blue: tactile left trial; dark blue: tactile right trial; orange: auditory left trial; red: auditory right trial. **b**, Example sessions of one TAs mouse (upper row) and one TAr mouse (lower row) showing licking patterns in sessions that switched from tactile to auditory task (left column) and from auditory to tactile task (right column). Each dot represents a single lick. Time was aligned to go cue onset. Optogenetic trials were excluded, while pre-licking trials were retained. Horizontal dashed line, the first transition trial. L, left trial; R, right trial. **c**, Behavioral performance of TAs mice during tactile-to-auditory or auditory-to-tactile transitions, mean ± s.e.m. across sessions (bootstrap, n = 42, 43 sessions respectively). Left and right trials were aligned to task switching separately. Pink, left trial; purple, right trial. Black ticks above the plot indicate trial windows with significant performance differences between left and right trials (*P*<0.05, two-sample t-test). Vertical dashed line, the start of task switching. Horizontal dashed line, chance level performance (50%). **d**, Same as c but for TAr mice (n = 63, 66 sessions respectively). **e**, Statistics on the number of trials for performance recovery after switching. Light green, TAs; dark green, TAr. \*\*\**P*<0.001, two-sample t-test. **f**, Schematic of optogenetic manipulation during the sample or delay epoch. **g**, Behavioral performance after left ALM inhibition. Thin line, single animal; thick line, average. \**P*<0.05, \*\**P*<0.01, \*\*\**P*<0.001, paired-sample t-test against control trials (TAs, 8 mice; TAr, 7 mice). **h**, Same as f for behavioral performance after right ALM inhibition.

We trained two cohorts of mice with distinct task switches (Fig. 1a). For the congruent switching, salient stimuli (large whisker vibrations, loud sounds 8Hz pure tone) signal right-choice trials in both tactile and auditory tasks, i.e. tactile and auditory tasks have a similar sensory-motor contingency (TAs). For the incongruent switching, tactile task retains salient-right and soft-left association, while auditory task reverses contingency (soft sound signals right-choice), i.e. tactile and auditory tasks have a reversed sensory-motor contingency (TAr). This design creates analogous rules for TAs mice (favoring cross-task generalization) versus conflicting rules for TAr mice (imposing computational interference). Critically, the tactile task itself (stimuli, timing, motor planning, and motor outputs) was identical across TAs and TAr cohorts, and the auditory was likewise identical across cohorts (except the sensory-motor mapping rule). This isolates the effect of rule alignment on transition behavior and on the compositionality, dimensionality and cross-task generalizability of population codes.

Mice performed either the tactile or the auditory task well (TAs mice: tactile task, 82.2 ± 6.5%; auditory task, 77.4 ± 5.8%. TAr mice: tactile task, 76.2 ± 5.5%; auditory task, 70.9 ± 4.4%. Mean ± s.d. across 85 sessions for TAs mice, 82 sessions for TAr mice), and transitioned between tactile and auditory tasks once or twice per session (Fig. 1b). Mice withheld licking well during the delay epoch (Fig. 1b), with low rates of pre-licking (Extended Data Fig. 1a. TAs mice: tactile task, 7.3 ± 10.7%; auditory task, 10.7 ± 13.7%. TAr mice: tactile task, 6.2 ± 7.3%; auditory task, 12.6 ± 12.4%. Mean ± s.d.) and no-response (Extended Data Fig. 1b. TAs mice: tactile task, 1.8 ± 2.9%; auditory task, 1.7 ± 2.8%. TAr mice: tactile task, 1.6 ± 1.9%; auditory task, 1.6 ± 2.2%. Mean ± s.d.). Intriguingly, TAs and TAr mice showed quite different characteristics during transition (Fig. 1c, d; see Extended Data Fig. 1c, d for single sessions; Lick latency did not fluctuate during task transition, Extended Data Fig. 1e – h). TAs mice maintained stable performance when transitioning modalities (Fig. 1c), reflecting efficient cross-task generalization. In contrast, TAr mice exhibited marked post-switch biases (e.g., impaired right-choice trials when switching tactile→auditory; Fig. 1d; see Discussion for transition mechanism). This asymmetry required TAr mice to undergo significantly more trials to achieve successful switching (Fig. 1e), highlighting how incongruent rules amplify behavioral conflict by impeding cross-task rule transfer.

To link rule structure to frontal cortical computation, we first optogenetically inhibited the anterior lateral motor cortex (ALM) during task performance (Fig. 1f, Methods). Unilateral inhibition during the sample epoch caused deficits in trials instructed by salient stimuli (right trials in both TAs and TAr tactile task, right trials in TAs auditory task, and left trials in TAr auditory task, Fig. 1g, h), implicating ALM’s role in sensory coding. Inhibition during the delay epoch induced an ipsilateral response bias, consistent with ALM’s role in motor planning^29,30,38^. These results position ALM as one critical node for sensorimotor transformation, and we thus recorded ALM population activity during task transition.

### Congruent and incongruent rules differentially shape shared neural representations

To investigate how rule congruence shapes neural coding representation, we recorded ALM ensemble activity in TAs and TAr mice during tactile and auditory tasks (Neuropixels 1.0 probes; Extended Data Fig. 2; Methods). Overall, we recorded 11,253 high-quality single units from ALM (TAs mice: 2,058 and 2,893 high-quality units in the left and right ALM from 6 mice with 44 and 43 recording sessions, respectively; TAr mice: 2,994 and 3,308 units in the left and right ALM from 8 mice with 59 and 57 sessions, respectively).

Putative pyramidal neurons were rigorously selected for further analysis (Methods). Neurons were classified based on selectivity for left or right choices during the sample, delay or response epoch in either tactile or auditory task. Neurons were further classified into one of nine categories (left-preferring, right-preferring and non-selective in tactile and/or auditory tasks; see example delay-epoch selective neurons in Fig. 2a, b; population activity patterns in Extended Data Fig. 3a-l).

**Fig. 2.**
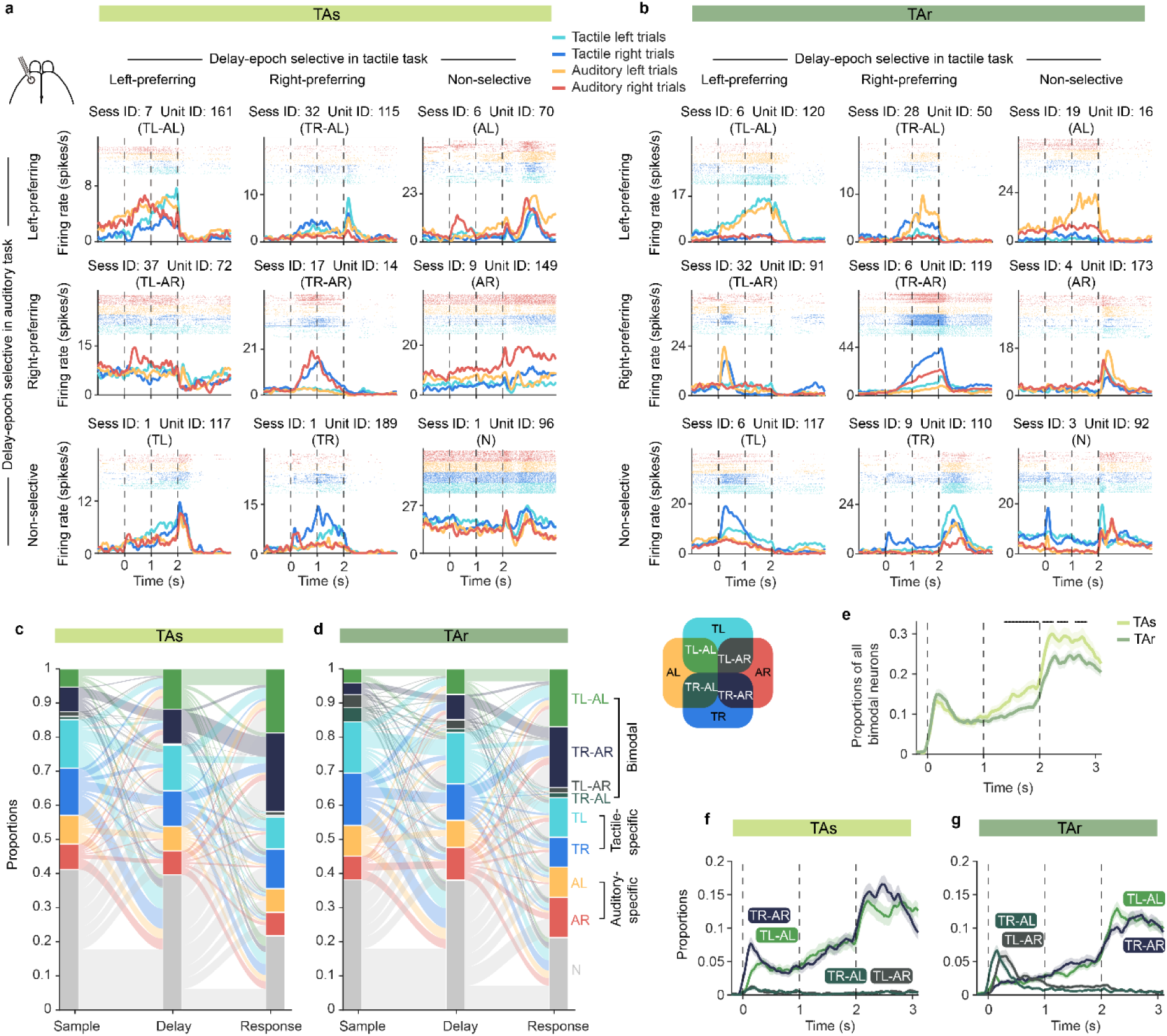
Activity patterns and shared representations in TAs and TAr mice. **a**, Example delay-selective neurons recorded in the left ALM from TAs mice. Neurons are arranged by delay-epoch selectivity in tactile task (horizontal axis) and in auditory task (vertical axis). For each neuron, session index and unit index are shown at top, with neuron type in the bracket. T, neuron shows selectivity in tactile task; A, neuron shows selectivity in auditory task; L, left-preferring; R, right-preferring; N, non-selective. Spike raster and trial-averaged peri-stimulus time histogram (PSTH) of correct trials are plotted for each neuron. PSTHs were calculated with a 1 ms time bin and averaged over 200 ms. Dashed lines, task epochs. **b**, Same as **a** but for example neurons from TAr mice. **c**, Sankey diagram to visualize changes of neuron selectivity over epochs in TAs mice. **d**, Same as **c** but for TAr mice. Venn diagram shows the definition of each selective neuron type. **e**, Changes in the proportion of all bimodal neurons over time. Mean ± s.e.m. (bootstrapping sessions 1,000 times). Black ticks, trial windows with significant difference (*P*<0.05, one-sided bootstrapping test). **f**, Proportions of each bimodal neuron-type in TAs. **g**, Same as **f** but for TAr.

In TAs mice with congruent rule, neurons frequently exhibited aligned selectivity across modalities (Fig. 2a, c): a neuron preferring right trials in the tactile task also favored right trials in the auditory task (Fig. 2a, unit ID 14, TR-AR neuron; see unit ID 161 for left-left preferring, TL-AL). These “bimodal” neurons (selective in both tasks) occupied a substantial fraction of ALM neurons, with their proportions increasing from delay to response epochs (Fig. 2c, e). This suggests that congruent rules promote shared coding schemas, enabling cross-task generalization through reusable neural ensembles.

In contrast, TAr mice with incongruent rule displayed a striking divergence: while bimodal neurons also increased over trial progression, the fraction is significantly lower compared with TAs (one-sided bootstrapping test; Fig. 2d, e), and significantly more neurons exhibited contradictory selectivity (e.g., right-preferring in tactile but left-preferring in auditory tasks; Fig. 2b, unit ID 50, TR-AL neuron; Fig. 2g; TAs versus TAr, *P* < 0.001, one-sided bootstrapping test during the sample and delay epoch). The early sample epoch featured transient populations with opposing tuning (TL-AR, TR-AL; Fig. 2g), reflecting unresolved conflict under reversed sensorimotor mappings. Over time, these contradictory neurons declined and were replaced by neurons with consistent task-specific selectivity (TL-AL, TR-AR; Fig. 2g). This temporal reconfiguration was absent in TAs mice (Fig. 2f), indicating that incongruent rules recruit more distinct, non-overlapping subpopulations for each task. The progression from conflicting to consistent bimodal tuning in TAr suggests that resolving rule conflict entails a gradual reallocation of neural resources within ALM.

### Congruent and incongruent rules differentially regulate cross-task generalizability and compositional neural representations

To quantify how rule congruence modulates cross-task neural representations, we analyzed ALM population dynamics using both cross-task decoding and targeted dimensionality reduction methods.

We first randomly selected a pseudo-population of neurons and trained a support vector machine (SVM) decoder (with 10-fold cross validation, see Methods) to differentiate sensory instruction (left vs. right trials instructed by stimuli), choice (left vs. right licking directions chosen by mice), and outcome (correct vs. error trials). The decoder reached high accuracy (∼80%) with a small group of randomly selected neurons (TAs tactile sensory, 100; auditory sensory, 250; tactile choice, 150; auditory choice, 250; tactile or auditory outcome, 50; TAr tactile or auditory sensory, 100; choice, 300; outcome, 50; Extended Data Fig. 4a, b). We then randomly selected 300 neurons to examine to which degree the decoder trained on one task can be generalized to the same task (within-condition decoder) or different task (cross-condition decoder, Extended Data Fig. 4c; Methods). For within-condition decoding, sensory information was greatest at the early sample epoch (Fig. 3a, b, left), whereas choice information gradually increased from sample to delay and reached a peak at the beginning of the response epoch (Fig. 3a, b, middle). Outcome information only emerged in the response epoch (Fig. 3a, b right), consistent with its definition. For cross-condition decoding, sensory information in the sample epoch barely generalized to the other task (Fig. 3a, b, left). In contrast, the choice and outcome information generalized well across tasks (Fig. 3a, b, middle and right). We quantified the cross-task generalizability and found that TAs and TAr mice showed marked difference. Decoders trained on tactile or auditory task variables achieved significantly lower cross-task generalization in TAr mice (Fig. 3c. The generalization index: TAs = 0.35 ± 0.18 vs. TAr = −0.17 ± 0.13 for sensory instruction; TAs = 0.80 ± 0.15 vs. TAr = 0.61 ± 0.16 for choice; TAs = 0.97 ± 0.04 vs. TAr = 0.91 ± 0.09 for action; TAs = 0.96 ± 0.03 vs. TAr = 0.90 ± 0.07 for outcome; Mean ± s.d.; *P* < 0.001, bootstrap test for equality of means; Methods), consistent with the contradictory selectivity patterns in TAr (Fig. 2).

**Fig. 3.**
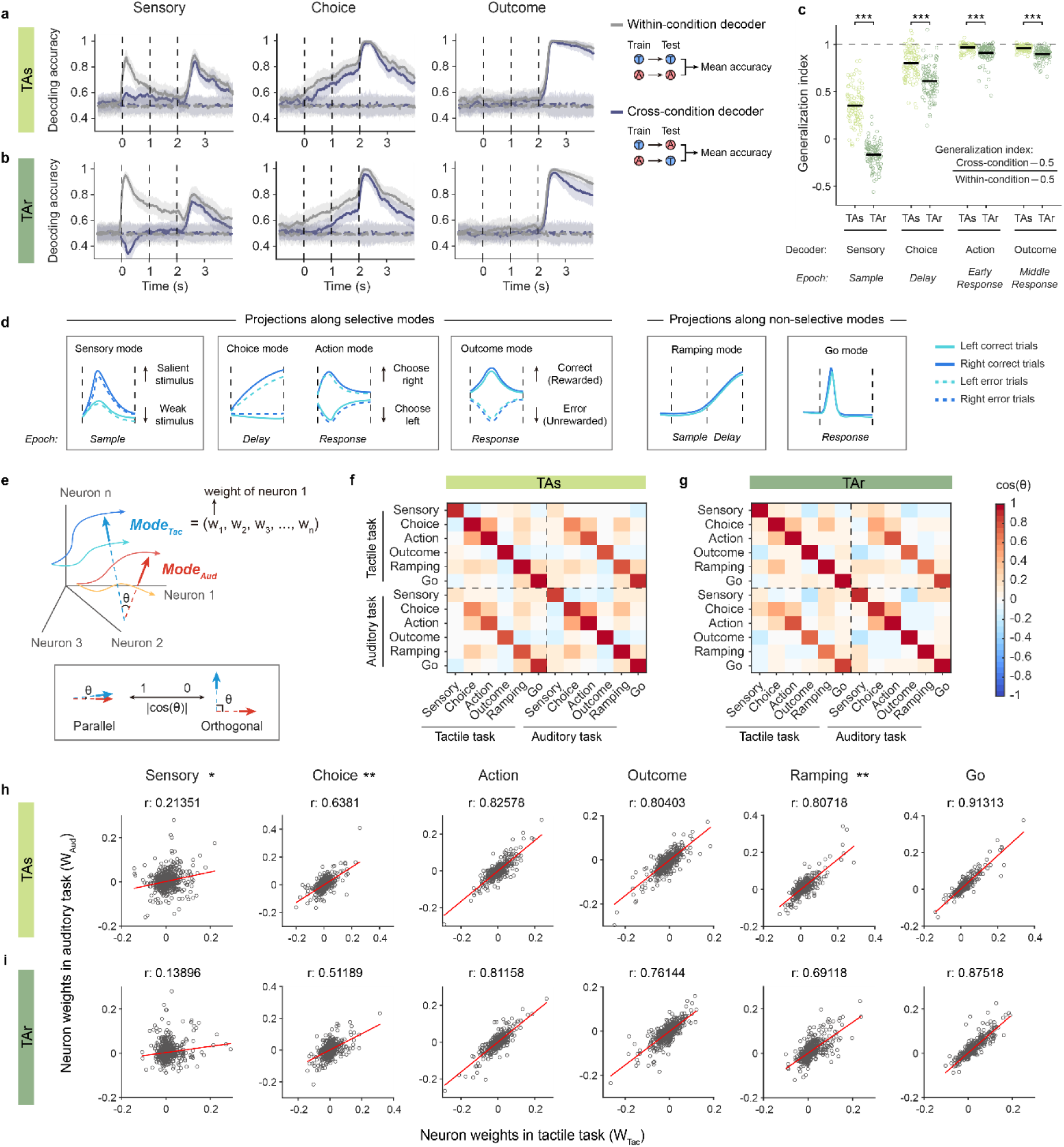
Cross-task generalizability and compositional coding in TAs and TAr mice. **a**, Within-condition and cross-condition decoding accuracy for sensory instruction, choice and outcome from recorded neurons in TAs mice, mean ± s.d. across100 runs. Solid line, decoder trained on true data; dashed line, decoder trained on label-shuffled data. **b**, Same as **a** but for TAr mice. **c**, Generalization index for each task variable. For sensory instruction and choice, generalization across tasks was higher in TAs group than in TAr group. Black bar, mean. Dashed line, complete generalization. \*\*\**P*<0.001, bootstrap test for equality of means. **d**–**e**, Schematics of activity modes and neural activity projections. **d**, Schematics of projections along tactile activity modes. Sensory, choice, action and response modes are selective modes, each specified by neural activity in task epochs (labeled below). Ramping and go modes are non-selective modes, capturing trial-type independent dynamics over time. **e**, Illustration showing neural population trajectories and angles between activity modes. In the *n*-dimensional neural state space, neural activity in each trial type evolves as a trajectory (thin arrow). Activity modes, labeled as ***Mode_Tac_*** and ***Mode_Aud_*** (thick arrows), are *n*-dimensional vectors with each entry representing neuronal weights. **f**, Cosine of angles between activity modes for TAs mice. Dashed lines separate tactile and auditory activity modes. **g**, Same as f for TAr mice. **h**, Correlation of neuron weights for corresponding tactile and auditory modes in TAs mice. Horizontal axis, neuron weights of tactile activity modes; vertical axis, neuron weights of auditory activity modes. Each circle indicates one neuron. Red line, linear fits. The r value at top denotes the Pearson correlation coefficient. **i**, Same as **f** but for TAr mice. For sensory, choice and ramping modes, the correlation coefficients were significantly different compared to those of TAs. \**P*<0.05, \*\**P*<0.01, one-sided permutation test.

To corroborate that task congruence regulates similarity of neural representations, we further conducted targeted dimensionality reduction that decomposed ALM activity into task-variable modes^19,39^ (Fig. 3d, Extended Data Fig. 6; Methods). Sensory, choice, action and outcome modes are selective modes that differentiate trial types in the sample, delay or response epochs. Ramping and go modes are non-selective activity modes, capturing the ramping activity during the delay epoch and the abrupt rise following the go cue, respectively. These activity modes capture a substantial proportion of activity variance (Extended Data Fig. 6) and are nearly orthogonal to each other within each task (Fig. 3f, g), showing as significant functional modules (Extended Data Fig. 7a, Extended Data Fig. 8a; *P* < 0.001, ePAIRS test; Methods). However, activity modes across tasks show distinct geometric relations, with the tactile and auditory sensory modes being roughly orthogonal, whereas other modes exhibit increasing degrees of parallelism (diagonal elements of the upper-right and lower-left matrix, Fig. 3f, g). We then visualized neuron weights for each activity mode in a *t*-SNE space (Extended Data Fig. 7b, Extended Data Fig. 8b; Methods). Neurons contributing to each activity mode were not randomly distributed, but rather exhibiting compositional modular patterns, with sensory modes largely segregated and choice/action modes roughly overlapping (Extended Data Fig. 7c, Extended Data Fig. 8c), consistent with compositional reuse.

Consistent with difference in cross-task decoding generalizability, TAs and TAr mice exhibited differential modulation in sensory, choice and ramping modes. We quantified cross-task alignment by correlating each neuron’s weights on matched modes across modalities (Fig. 3h, i). Alignment increased from sensory and choice to action, outcome, ramping, and go modes, indicating greater sharing for late-stage variables. However, alignments for sensory, choice and ramping were significantly weaker in TAr (sensory *P* = 0.0336, choice *P* = 0.002, ramping *P* = 0.0063 vs. TAs; one-sided permutation test, Methods; Fig. 3h, i), as neurons contributing to tactile and auditory representations occupied distinct subspaces (see t-SNE visualization of non-overlapping neural subpopulations, Extended Data Fig. 7c, 8c). Consistently, sensory, choice, and ramping modes were less parallel in TAr (*P* = 0.0306, 0.002, 0.0068; Fig. 3f, g). In contrast, TAs mice leveraged more shared subspaces for sensory-to-action transformations (Fig. 3h; Extended Data Fig. 7c), consistent with the higher generalizability across tasks. Together, these analyses show that congruent rules promote efficient compositional reuse of ALM ensembles across tasks, whereas incongruent rules demand more distinct subpopulations, leading to reduced cross-task generalizability.

### Incongruent rules recruit a larger fraction of nonlinear mixed selectivity (NMS) neurons

Neurons in ALM exhibited diverse, epoch-dependent selectivity that reorganized after task switches. We asked whether incongruent rules preferentially engage NMS neurons, with responses depending on interactions between task identity (modality) and other variables. For each neuron and epoch (sample, delay, response), we fitted linear models with terms for modality, sensory, choice and outcome, plus their interactions (Methods). Most neurons are fitted successfully with significant models (*F*-test, *P* < 0.05; the sample epoch, 887/1125 and 1293/1575 in left ALMs of TAs and TAr, respectively, 1119/1466 and 1364/1722 in right ALMs of TAs and TAr; the delay epoch, 926/1125, 1289/1575, 1142/1466, 1359/1722; the response epoch, 1031/1125, 1443/1575, 1339/1466, 1558/1722). Neurons are further classified into four categories based on their specificity of responses: NMS (encoding interactive variable combinations, see example neurons in Fig. 4a–d, Methods), linear mixed selective (LMS; additive coding, see example neurons in Extended Data Fig. 9a–d), pure selective (PS; single variable coding), or non-selective (NS).

**Fig. 4.**
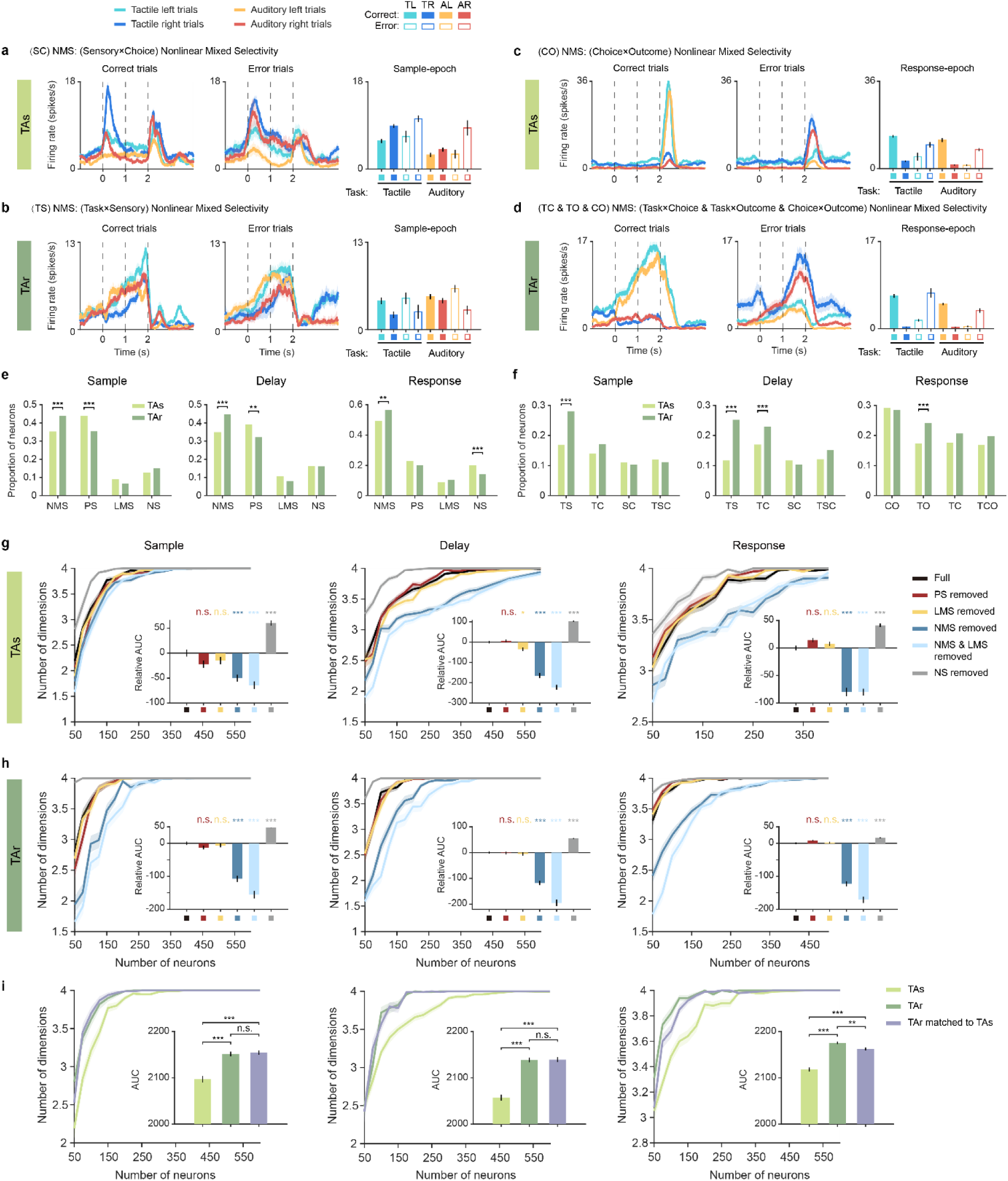
Incongruent task rules relates to higher nonlinear mixed selectivity. **a-b**, Example NMS neurons fitted sample-epoch firing rate with task modality (T), sensory stimulus (S) and choice (C). Left panel, PSTHs of correct trials; middle panel, PSTHs of error trials; right panel, averaged sample-epoch firing rate. Mean ± s.e.m. **a**, an example NMS neuron of TAs showing nonlinear mixed selectivity to sensory stimulus and choice (Sensory×Choice, SC). **b**, an example NMS neuron of TAr showing nonlinear mixed selectivity to task modality and sensory stimulus (Task×Sensory, TS). **c–d**, Example NMS neurons fitted response-epoch firing rate with task modality (T), choice (C) and outcome (O). **e**, Proportions of NMS, PS, LMS and NS neurons across different epochs. **f**, Proportions of NMS neurons showing each selectivity-type across epochs. Neuronal activities were fitted with T, S and C during sample and delay epochs, and with T, C and O during the response epoch. *P* values, two-sided permutation test. **g**–**h**, Effects of removing different neuron types on the dimensionality of neural representations. The number of dimensions gradually increased with the number of neurons. For each condition, proportions of (remained) neuron types were preserved in each iteration. Mean ± s.e. across 100 runs. The inset shows the quantification of the relative effects of neuron type removal. *P* values obtained by comparing the area under curve (AUC) of the neuron-removed group with that of the full population (black), two-sided permutation test. **i**, Matching neuron type proportions of TAr dataset to TAs dataset. Mean ± s.e. across 100 runs. *P* values, two-sided permutation test. In all panels, \**P* < 0.05, \*\**P* < 0.01, \*\*\**P* < 0.001, with Bonferroni correction for multiple comparison.

In contrast to TAs mice, TAr mice exhibited an overrepresentation of NMS neurons across all epochs, and a lower fraction of PS neurons (Fig. 4e, Extended Data Fig. 9e, f). Specifically, TAr had significantly more Task×Stimulus (TS) NMS neurons than TAs during the sample and delay epochs (Fig. 4f, left ALM, TAs vs. TAr *P* < 10^4^; see Extended Data Fig. 9g for the right ALM, sample epoch TAs vs. TAr *P* = 0.0054, delay epoch TAs vs. TAr *P* < 10^4^; two-sided permutation test with Bonferroni correction for multiple comparisons). During the delay epoch when choice information ramped up (Fig. 3a, b), TAr in addition recruited Task×Choice (TC) NMS neurons that combined task modality and choice (Fig. 4f, left ALM, TAs vs. TAr *P* = 0.001; see Extended Data Fig. 9g for the right ALM, TAs vs. TAr *P* < 10^4^). In the response epoch when the outcome information first emerged (Fig. 3a, b), TAr recruited significantly more Task×Outcome (TO) neurons that combined task modality and outcome (Fig. 4f, left ALM, TAs vs. TAr *P* < 10^4^; see Extended Data Fig. 9g for the right ALM, TAs vs. TAr, *P* = 0.0012). Thus, incongruent rules systematically increase the reliance on nonlinear mixed selectivity, recruiting neurons that conjunctively encode task identity with stimulus, choice, and outcome at their respective epochs.

### Incongruent rules shape geometry and dimensionality of neural representations

Previous studies have indicated that nonlinear mixed selectivity increases the dimensionality of neural representations^5,9–11^. Given the elevated NMS recruitment in TAr mice and their transition costs under reversed sensorimotor contingencies (Fig. 1c–e), we hypothesized that incongruent rules demand higher-dimensional codes to segregate conflicting sensorimotor mappings, whereas congruent rules reuse shared, low-dimensional subspaces for fast task transition (Fig. 5a, left).

**Fig. 5.**
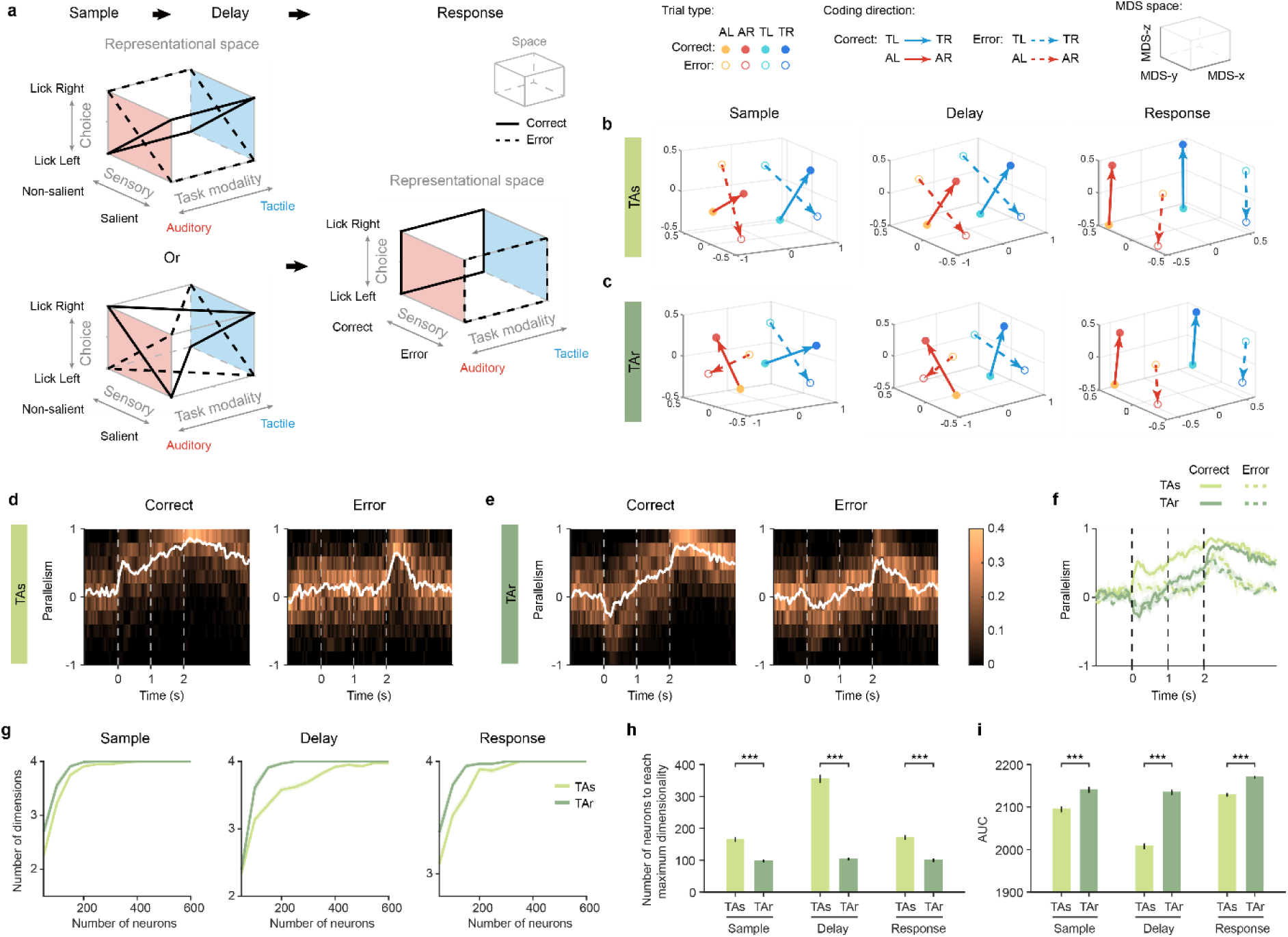
Task rules shape geometry of neural representation. **a**, Schematic of neural representations of task variables. **b**–**c**, Embeddings of mean neural activity during the sample, delay and response epochs in 3D MDS spaces. Each circle represents trial-averaged population activity for a specific trial type. Filled circles, correct trials; open circles, error trials. Solid arrows, coding directions (CDs) for correct trials; dashed arrows, CDs for error trials. The axes of the MDS spaces are rotated so that embedding of TAs and TAr in different epochs are comparable. **d**–**e**, Distribution of angles between tactile and auditory CDs. Here CDs were calculated for single sessions. Left, cosine values of the angles between correct tactile and auditory CDs; right, cosine value of angles between error tactile and auditory CDs. Cosine similarity quantifies the parallelism between CDs. White line, median across sessions. **f**, Overlay of the median values shown in d– e, median ± standard error of median (bootstrap) across sessions. Solid line, values of correct CDs; dashed line, values of error CDs. **g**, Number of representational dimensions as the number of neurons increased. Mean ± s.e. across runs. **h**, Quantifications of the number of neurons required to reach maximum dimensions. Mean ± s.e. across runs. **i**, Quantifications of area under curves (AUCs) in g. Mean ± s.e. across runs. \*\*\**P* < 0.001, two-sided permutation test, Bonferroni correction for multiple comparison.

To directly visualize geometric organizations, we embedded population activity into a 3D space using multidimensional scaling (MDS; Methods). MDS embedding of trial-averaged activity (correct or error trials for left-or right-trials instructed by sensory stimuli) revealed striking differences in coding directions (CDs; vectors separating left-vs. right-trials, shown as arrows in Fig. 5b, c) across rule contexts. In TAr mice, tactile and auditory CDs were nearly orthogonal during the sample and delay epochs (Fig. 5c), indicating modality-specific subspaces that minimize interference between reversed contingencies (e.g., left = “soft” in tactile vs. right = “soft” in auditory). In TAs, CDs were largely parallel across modalities (Fig. 5b), consistent with shared coding axes under congruent rules. During the response epoch, CDs converged in both contingencies (Fig. 5b, c), reflecting common motor outputs after the go cue (e.g. licking left or right water spouts) and outcome (Fig. 5a, right). We quantified the parallelism of tactile and auditory CDs by calculating the cosine angle (Fig. 5d, e). A value near 0 during early epochs confirmed the orthogonality of CDs in TAr (Fig. 5e). Similar observations held for error trials (Fig. 5b–e), although the differences in parallelism were considerably smaller (Fig. 5f).

We further estimated the dimensionality of neural representations under different rule contexts (Methods). The number of dimensions gradually increased with the number of neurons until reaching a plateau (Fig. 5g). TAr mice required significantly fewer neurons to reach the plateau across all epochs (Fig. 5h; TAs vs TAr, *P* < 0.001 for all epochs and both hemispheres; also see Fig. 5i for comparison of area under curve; Extended Data Fig. 10a – c for right ALM). This lower neuron requirement implies that TAr populations encode information in a higher intrinsic dimensionality, consistent with the orthogonalized CDs observed in MDS analysis (Fig. 5c).

To isolate contributions of different neuron classes to representations, we recomputed the dimensionality after selectively removing NMS, LMS, or PS neurons (Fig. 4g, h for left ALM; Extended Data Fig. 10d, e for right ALM; Methods). Removing NMS neurons markedly reduced dimensionality in both contingencies, increasing the neurons required to reach plateau (dark blue line; *P* < 0.001, permutation test, Fig. 4g, h). Removing PS neurons, which had comparable proportions to NMS neurons in the sample and delay epochs (Fig. 4e), did not reduce the dimensionality in TAr mice (red line, Fig. 4h). Interestingly, although removing LMS neurons alone had minimal impact (yellow line), likely due to lower prevalence, removing all mixed selectivity (NMS and LMS) further collapsed dimensionality (light blue). These results suggest that the mixed selectivity, particularly the nonlinear component, is critical for the high-dimensional representation. Notably, matching the proportions of NMS, PS, and LMS neurons in TAr to those in TAs did not eliminate the dimensionality gap (Fig. 4i, Extended Data Fig. 10f; Methods), indicating that NMS contributes to dimensionality beyond mere numerical abundance.

### Reorganization of neural representations predicts task transitions

One intriguing phenomenon is the markedly different behavioral performance of TAs and TAr mice during task transitions. TAs mice maintained stable performance, whereas TAr mice exhibited pronounced, rule-specific choice biases (Fig. 6a, d). Two competing hypotheses can potentially explain the transition difference. First, new sensory inputs were blocked or gated in ALM, and sensory information only recovered gradually post-switch. Second, ALM neurons encode new sensory stimuli but failed to link them to appropriate choices and actions.

**Fig. 6.**
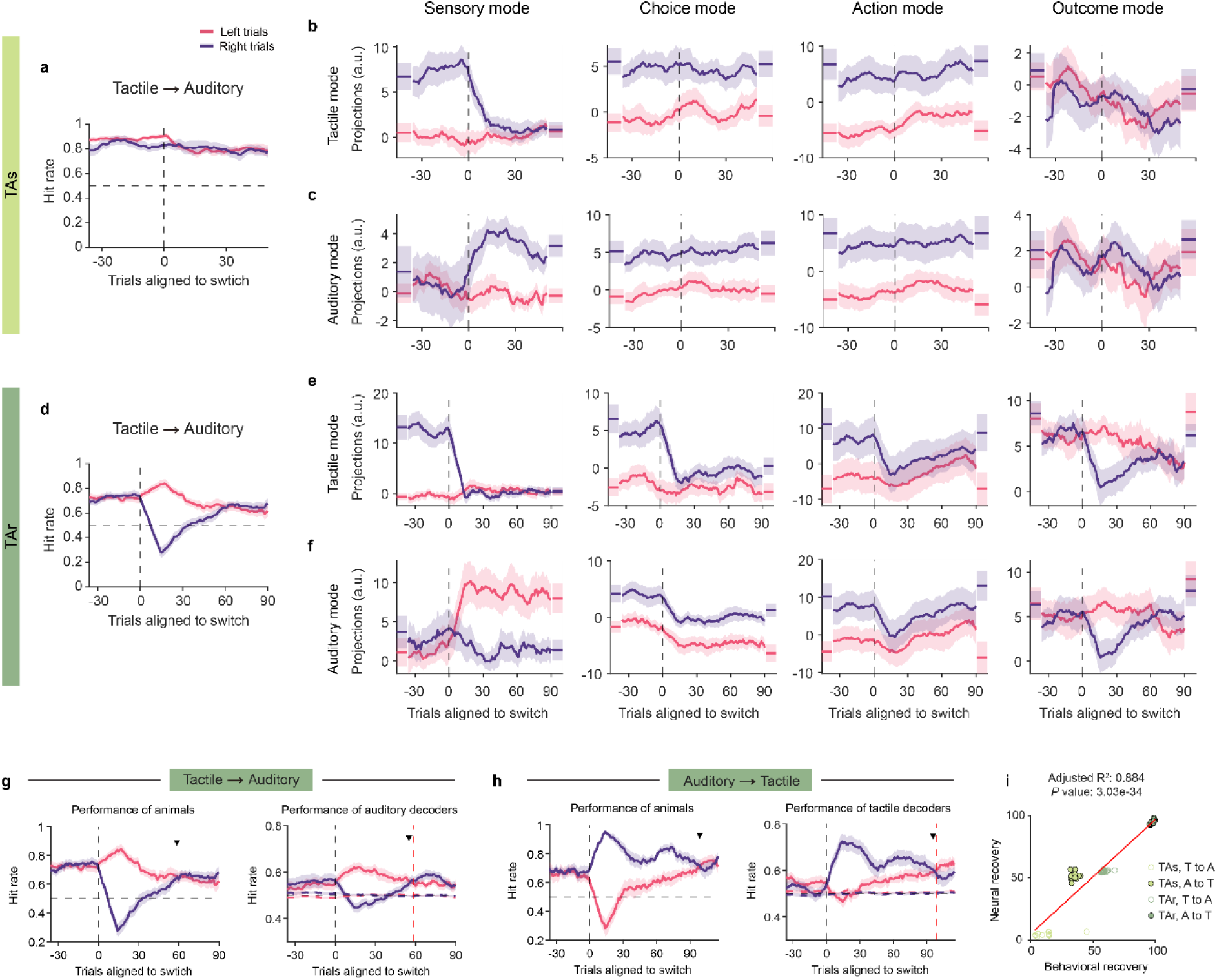
Reorganization of neural representations predicts task transitions. **a**, Behavioral performance of TAs mice. Same as Fig. 1c but 14 sessions with electrophysiology are included. Left and right trials were grouped according to sensory instructions in all panels. **b**, Projections of single-trial activity along sensory, choice, action and outcome modes defined from the tactile task. Mean ± s.e.m. (bootstrap) across sessions. Vertical dashed line, task switching. Short dashes on the left and right sides, projections of stable trials from tasks before and after switching. **c**, Same as **b**, but for projections along auditory activity modes. **d**–**f**, Same as a-c but for TAr mice with 24 sessions included. **g**, Behavioral performance (left) and auditory neural activity predicted performance (right) during tactile-to-auditory task switches in TAr mice. Mean ± s.e.m. (bootstrap) across sessions. Downward triangle marks the trial on which performance of left and right trials achieved the same level. Red dashed line in the right panel, trial of behavioral recovery. Dashed pink and purple lines in the right panel, performance of decoders with trial labels shuffled. **h**, Behavioral performance (left) and tactile neural activity predicted performance (right) during auditory-to-tactile task transitions in TAr mice. **i**, Correlation between behavioral recovery and neural recovery, calculated from decoder performance of the post-transition task. Red line, linear regression.

To disentangle these hypotheses, we applied targeted dimensionality reduction to extract activity modes (sensory, choice, action, outcome) from stable trials. We then projected single-trial activity during transitions onto these fixed modes (Methods). This approach enables analysis of the dynamic coding of behavioral variables during the unstable transition period. Tactile-to-auditory and auditory-to-tactile transitions were separated for analysis. During tactile to auditory transitions, projections along auditory sensory mode rose immediately after the switch in both TAs and TAr, while those along tactile sensory mode collapsed (Fig. 6b, c, e, f, left panels), ruling out sensory gating in ALM. Analogous results held for auditory→tactile transitions (Extended Data Fig. 11a–f).

In TAs, choice and action projections remained aligned with their pre-switch structure, matching stable behavior (Fig. 6b, c, middle panels). In TAr, right-trial activity collapsed toward left-trial projections along both choice and action modes (Fig. 6e, f, middle panels), mirroring the leftward behavioral bias. Outcome-mode projections for right trials deviated from pre-switch baseline, indicating maladaptive reward association with erroneous left choices (Fig. 6e, f, right panels). Analogous results held for auditory to tactile transitions (Extended Data Fig. 11a–f). Thus, although ALM rapidly encodes the new sensory stimuli in both contingencies, under incongruent rules it initially fails to remap sensory evidence onto the correct choice/action axes, supporting the hypothesis of sensorimotor mapping failure.

To further investigate how neural representations of choice relate to animals’ behaviors, we analyzed the transition process based on decoding analyses. We trained single-trial choice decoders on stable trials of the post-switch task (Methods). We evaluated the decoders on transition trials grouped by sensory instruction, rather than choice, treating the decoders as agents performing the task. This procedure ensured that behavioral and neural dynamics during task transitions were directly comparable (Fig. 6g, h). Decoder performance recapitulated the asymmetric divergence between left and right trials after switches. We then quantified recovery as the trial window in which left and right performance converged. Recovery times for behavior and neural decoders were strongly correlated across conditions (Fig. 6i), showing that the reorganization of ALM choice representations predicts the time course of behavioral stabilization.

## Discussion

Investigating how the brain supports multiple cognitive tasks offers insight into cross-task generalization flexibility of the neural system. In our behavioral paradigms, both TAs and TAr involved tactile and auditory tasks, but were designed with distinct sensorimotor mapping rules (congruent vs incongruent). By analyzing ALM neural activity between tasks, we observed the diversity of neuronal activities and the growing number of rule-dependent bimodal neurons over epochs (Fig. 2). By dissecting neural activity, our results uncovered congruence-dependent modular and compositional representations of task variables, and related these neural representations to cross-task generalizability (Fig. 3).

Our study further experimentally demonstrated that congruent and incongruent rules influence the mixed selectivity at single neuron level (Fig. 4) and the representational geometry and intrinsic dimensionality at population level (Fig. 5), which may in turn affect the cross-task generalizability of information and the requirement of cognitive resource during task transition. ALM rapidly encoded new sensory stimuli after task switch, nevertheless TAr mice initially failed to remap these inputs onto appropriate choice representations. The recovery of choice representations predicted behavioral recovery, underscoring ALM’s role in reconfiguring sensorimotor mappings (Fig. 6). By integrating task rules, mixed selectivity, representational dimensionality, and transition mechanisms, our study provides a coherent perspective on the neural principles that enable flexible and adaptive behaviors across tasks. These insights may also inspire AI systems, suggesting strategies for multi-task learning, modular network design, and adaptive decision-making in changing contexts.

### Compositional coding, nonlinear mixed selectivity, and cross-task generalization

Previous studies have shown that neural representation can be either generalist or specialist to perform different tasks^40–44^. Abstract knowledge, such as orientation^23^, frequency^22^, and spatial location^26^, as well as cognitive variables such as decision confidence^24^, can be shared across sensory modalities. Without factorizing tasks into constituent variables, cross-modal generalization of neural representations can be difficult to precisely estimate. Our study decomposed each task into several components encoding sensory perception, decision making, motor execution, and reward consumption. In ALM, generalization increased over time across tasks, consistent with its premotor role and convergent motor outputs. Sensory information barely generalized in ALM, in line with its failure to abstract contextual cues despite its necessity for flexible stimulus-response remapping^37^. Whether stimulus salience or intensity is represented abstractly elsewhere to support task-structure inference, and how such codes transform across cortical modules, remains open. Frontal and parietal cortices are candidates for mediating cross-modal generalization and may provide top-down constraints that align or separate subspaces in ALM depending on rule congruence. Future causal and multi-area recordings could test how these networks transform sensory evidence into flexible motor plans and govern rule generalization under congruent versus incongruent sensorimotor mappings.

Recent work shows that ALM generated new preparatory activity when mice learned novel task contexts involving reversed contingency or changed stimuli, supporting the notion that these motor memories were stored in combination with specific contexts^28^. While such context-specific representations may prevent interference, they can compromise flexibility^44^. Here in well-trained animals, we observed contingency-dependent cross-task generalization of choice and non-selective ramping activity, consistent with the notion that choice and ramping modes are encoded by overlapping neural populations^19^. How compositional codes emerge with learning, and whether simultaneous versus sequential training differentially sculpts subspace alignment and mixed selectivity, are important next steps.

In the TAr paradigm with incongruent rules, increased NMS neurons and a higher dimensionality across epochs indicated twisted sensorimotor mappings. The increased dimensionality allows more flexible readout of information by downstream areas, but it can deteriorate cross-task generalization of neural representations, explaining slower transitions in TAr mice. Importantly, matching the proportions of NMS, LMS, PS and NS neurons did not abolish the dimensionality gap, suggesting an intrinsically high-dimensional coding property of the neuronal population. While abstract representations of task variables can facilitate generalization, they retained a relatively higher dimensionality than a fully factorized representation^45^. This trade-off between dimensionality and generalization may be essential for maintaining both stability and flexibility in neural systems^46,47^.

### Mechanisms of Task Transition

By projecting activity in transition trials onto sensory and choice coding directions, we ruled out sensory gating as ALM encoded new sensory modality immediately after task switch. Indeed, ALM neurons can faithfully encode sensory information even for distractors^32^. Instead, the collapse of choice decoding in TAr indicates a remapping failure that sensory information was not linked to the appropriate choice. Notably, the representations of left and right choices stabilize at different rates under TAr (Extended Fig. 11h, j). Following tactile-to-auditory switching, the auditory choice decoder recovered faster for left than right trials (Extended Fig. 11h). After auditory-to-tactile switching, the tactile decoder recovered better for right than left trials (Extended Data Fig. 11j). Thus, under incongruent task rules, TAr mice preferentially engaged choice mappings for the more salient stimulus-outcome mapping, paralleling behavioral biases during transitions. This asymmetry did not arise from absent sensory drive to cortex as both salient and non-salient stimuli evoked robust neural responses in the primary sensory cortices (Extended Fig. 12). In TAs mice, the parallel coding directions and higher cross-task generalizability ensured a smoother remapping, and the choice decoding of left and right trials recovered more synchronously (Extended Data Fig. 11g, i). Together, these results implied that task transitions depend jointly on stimuli salience and task rules. The unequal salience of left and right trials, unlike in other tasks such as categorization or stimulus comparison where choices are nearly balanced^24,43,48^, highlights mechanisms of choice formation under highly informative sensory inputs. Future work involving continuous stimuli can further enhance understanding of cross-task representation reorganization underlying flexible sensorimotor reassignment.

Prior studies of task transitions often used context-dependent paradigms with identical sensory inputs but rule changes, requiring attention shift across stimulus features (e.g., shape or color)^49^ or sensory modalities^36^. These studies highlighted the role of task belief and pre-stimulus neural states in guiding switches. In our paradigm, animals typically experienced one or two transitions per session. How context and block size influence task transition dynamics remains an open question.

In our data, the recovery of choice-related activity in ALM closely tracked behavioral recovery, and slightly preceded it under incongruent rules (Fig. 6g, h). The timing difference is not observed in a previous study where mice switched between three stimulus-response sets^50^. Several design differences may account for this discrepancy. First, our tasks included a delay epoch that required animals to maintain choice, providing a clear time window to examine choice formation. Second, choices were not consistently symmetric across tasks in the previous study, making the choice dynamics difficult to compare. Nevertheless, our analysis relied on session-averaged traces, as performance of both animals and choice decoders in individual sessions was noisy, obscuring clean recovery estimates. Future experiments with larger, simultaneously recorded neuronal populations per session should sharpen single-session readouts and may uncover how nonlinear mixed selectivity contributes to task transitions.

## Acknowledgements

We thank Yang Zhou and Quan Wen for comments on the manuscript. This work was funded by the Ministry of Science and Technology of China (2021ZD0203600); the National Natural Science Foundation of China (32525032, 32170998 and 32021002); and the support from THU-IDG/McGovern Open Laboratory of Shared Instruments for Brain Science.

## Author contributions

X.C. and Z.V.G. conceived and designed the experiments. X.C. performed the experiments with help from X.Y. and X.X.Y. X.X.Y. helped with behavioral apparatus setup and project design. X.Y. helped with mice training and electrophysiology. X.C. analyzed data with comments from other authors. X.C. and Z.V.G. wrote the paper with inputs from all authors.

## Data and code availability

Data and the codes generated during this study will be made available prior to publication of the manuscript. Any additional information required to reanalyze the data reported in this paper is available upon reasonable request.

## Author Information

Correspondence and requests for materials should be addressed to.

## Methods

### Mice

This study was based on data from seventeen mice (age > postnatal day 60, male). PV-ires-cre mice (JAX 008069)^51^ crossed with Rosa26-LSL-ReaChR-mCitrine mice (JAX 026294)^52^ were used for photoinhibition and electrophysiology.

All procedures were approved by the Institutional Animal Care and Use Committee (IACUC) of Tsinghua University. Mice were maintained on a 12/12 reversed light/dark cycle, 22–26℃ with water ad libitum, and behaviorally tested during the dark phase. Mice were housed individually after craniotomy to prevent injury. All surgery procedures were performed aseptically under 1.5–2% isoflurane (R510-22-10, RWD) anaesthesia. Flunixin meglumine (Sichuan Dingjian Animal Medicine Co., Ltd) was injected subcutaneously (1.25 mg/kg) during surgery and postoperatively for at least three days to reduce inflammation. After the surgery, mice were allowed to recover for at least three days with free access to food before food restriction. A typical behavioral session lasted 1–2 hours and mice obtained milk as reward during the behavior. Mice received supplementary pellet food after training to maintain a stable body weight, typically 1.8–3 g depending on their performance. On days not tested, mice received 3–4 g pellet food.

### Surgery

Mice were prepared with a clear-skull cap and a head-bar^29^. The scalp and periosteum over the dorsal skull were removed. A thin layer of cyanoacrylate adhesive (Krazy glue, Elmer’s Products Inc.) was directly applied to the exposed skull. A custom made titanium head-bar was glued on the skull over cerebellum with its anterior edge aligned with the suture lambda. A layer of clear dental acrylic (#1223-clear, Lang Dental Jet Repair Acrylic) was applied over the cyanoacrylate adhesive and head-bar to cover the intact skull. A heated pad (Harvard Apparatus) was used to maintain the mice’s body temperature during operations and postoperative recovery.

### Behavior

The behavioral paradigms are designed based on the delayed-response task^29,38^. Head-fixed mice were trained to choose lick spouts instructed by stimuli from different sensory modalities. Tactile and auditory tasks involve distinct sensory inputs but identical motor outputs (licking the left or right port).

Before training, mice were food restricted and given 2 g of pellet food and 3–5 ml of 20% milk (w/v, Pet-Ag Inc.) per day. Mice were then acclimated to head fixation with free access to milk via two spouts (4.4 mm between spouts) for 2–3 days. Each lick received 2–4 μL reward and was accompanied by a 5 kHz auditory go cue (0.1 s, 70 dB).

Mice were then randomly allotted to two groups: one learning the TAs paradigm and the other learning the TAr paradigm. Mice began learning the tactile and auditory tasks on the same day, with 1–2 transitions between tasks each day. Mice withheld their licking during the delay epoch and chose spouts after the go cue, which indicated the start of the response epoch, during which correct licks were rewarded with 2 μL milk, while error licks triggered a 1-s timeout. Pre-lickings before the response epoch also triggered a prolonged delay epoch (1-s timeout).

For tactile stimuli, a mental pole was placed on each side of the mouse (1.4 cm between poles and equal distance from the whisker pads), with each pole attached to the shaft of a mirror galvanometer. The intensity and frequency of the whisker vibration was controlled by the current input from myRIO (National Instruments, myRIO-1900) to the galvanometer. A peak velocity of 408°/s was set for soft vibration (tactile left trials), and 1838°/s for strong vibration (tactile right trials) in both TAs and TAr mice’s tactile tasks.

For auditory stimuli, a speaker (Audax TW025A20 1” Titanium Dome Tweeter) was positioned approximately 20 cm in front of the head-fixed mice. The frequency, decibel level, and number of clicks for the pure tones were adjusted by MATLAB (MathWorks) parameters and output through myRIO. Pure tones at 65–70 dB (8 kHz, 10 clicks/s) were used for high-decibel stimuli, while 45–50 dB (8 kHz, 10 clicks/s) were used for low-decibel stimuli. For TAs mice, low-decibel stimuli indicate left trials, while high-decibel stimuli indicate right trials. In contrast, the contingency is reversed for TAr mice: low-and high-decibel stimuli instruct right and left trials, respectively. Another speaker was placed beneath this speaker to emit the 5 kHz go cue.

### Photoinhibition

Before optogenetic manipulation, the dental acrylic over the skull was polished and covered by a thin layer of clear nail polish (#72180, Electron Microscopy Sciences). To inhibit ALM, light from a 594 nm laser (Obis LS, Coherent) was controlled by an acoustic-optic modulator (AOM; MTS110-A3-VIS, Quanta Tech; extinction ratio 1:2000) and a mechanical shutter (Uniblitz LS6S2T0, Vincent Associates). Light was delivered to either the left or right ALM (anterior–posterior, AP: 2.5 mm; medial–lateral, ML: ±1.5 mm; relative to bregma) through the clear-skull cap implant using a 2D scanning galvo system (GVS012, Thorlabs).

We employed 40-Hz photostimulation with a sinusoidal temporal profile and a mean laser power of 1.5 mW. The photostimulation lasted for 1 s, including a linear ramp-down for 200 ms to minimize the rebound excitation after photoinhibition. Light was randomly delivered during the sample or delay epoch in 25% of trials. There was no photostimulation during task transitions. To prevent mice from distinguishing optogenetic trials from control trials using visual cues, a masking flash (1 ms pulses, 10 Hz) was delivered using 470-nm and 590-nm light-emitting diodes (Luxeon Star) throughout the trial (from −1 to 3 s relative to sample onset).

### Electrophysiology

Recordings were conducted through a craniotomy made 1–2 days before the recording. The craniotomies were centered over the left and right ALM (AP: 2.5 mm; ML: ±1.5 mm, relative to bregma) with a diameter of 0.5–1.0 mm. Extracellular spikes were recorded acutely using Neuropixels 1.0 probes. Probes were slowly inserted into the left and right ALM with an angle perpendicular to the cortex (±14° relative to the vertical, Extended Data Fig. 2a left). Recording depth was inferred from manipulator (MPC200, Sutter Instrument) readings. After completion of probe insertion, a drop of silicone gel (Part A, 3-4680 Silicone Gel Clear-Blue, DOWSIL) was applied over the craniotomy. The brain tissue was allowed to settle for several minutes before the recording started. After recording, the tissue was covered by silicone gel as artificial dura (Part A:Part B, 1:1) and the craniotomy was then covered by Kwik-Sil (World Precision Instruments).

Spikes from two simultaneously recorded probes were acquired at 30 kHz using the PXIe Acquisition Module and SpikeGLX 3.0 software (https://billkarsh.github.io/SpikeGLX/). A synchronization pulse (1 Hz) was sent to both the head-stage and the National Instrument DAQ card (NI PXIe-6341). Behavioral bitcode on the analog channel and the sync waves were coordinated through NI DAQ card. Signals from the PXIe Acquisition Module and the NI DAQ card were synchronized using the sync waves.

### Reconstruction of probe locations

To confirm the recording locations, CM-DiI (#C7001, Invitrogen) was applied to the tip of the Neuropixels probe to label the recording tracks. After the last recording session, mice were perfused transcardially with PBS followed by 4% paraformaldehyde (PFA). The brains were fixed overnight in 4% PFA and then cleared with uDISCO^53^. We imaged the cleared brain with Zeiss Lightsheet Z.1. Signals from the 488-nm and 638-nm channels were collected. The reconstruction was done using brainreg (https://github.com/brainglobe/brainreg) The signals from the 638-nm channel were used to register the mouse brain to the Allen Brain Reference Atlas. The obtained parameters were then applied to signals from the 488-nm channel to reconstruct the probe tracks (Extended Data Fig. 2a right).

### Behavioral data analyses

To analyze performance during task transitions, we excluded optogenetic trials and no-response trials. Hit rate was calculated as the proportion of correct trials using a sliding window of either 20 (Fig. 1) or 15 (Fig. 6, Extended Data Fig. 11) trials. Significant difference between left-and right-trial performance was assessed using two-sample *t*-test. A black tick was plotted above the middle trial window when three consecutive trial windows showed significance (Fig. 1c–d and Extended Data Fig. 1e–f).

To quantify the behavioral effects of photoinhibition, we excluded pre-licking trials across all conditions (control and photoinhibition). For each mouse, sessions were pooled to ensure a sufficient number of optogenetic trials for analysis. Hit rate was then calculated as the proportion of correct trials. Significance of the performance change in each photoinhibition condition was determined using paired-sample t-test.

### Electrophysiological data analyses

#### Neuropixels probe recording preprocessing

The extracellular recording traces were first band-pass filtered (300 Hz–10 kHz) and corrected (artifacts removed) using CatGT. We used an open-source pipeline for preprocessing SpikeGLX data, spike sorting and quality control (https://github.com/jenniferColonell/ ecephys_spike_sorting). Spike sorting was done using Kilosort2.0 (GitHub - jenniferColonell/KS20_for_preprocessed_data: Release version of KS2 for preprocessed data) or Kilosort3 (GitHub - MouseLand/Kilosort: Fast spike sorting with drift correction for up to a thousand channels). Units were selected based on the following quality metrics criteria: cluster_KSLabel: good; cluster_group: good; firing_rate > 0.1 Hz; amplitude_cutoff < 0.1; and isi_violations < 0.5. ALM units were obtained from a depth range of 0 to 1200 μm.

Spike duration was computed as the trough-to-peak width of the mean spike waveform. Units with mean firing rate > 1 spike/s and duration < 0.9 ms were included for further classification (Extended Data Fig. 2b–e). Units with duration < 0.25 ms were classified as putative fast-spiking neurons (FSs, 591 of 6,784) and units with duration > 0.3 ms as putative pyramidal neurons (PNs, 5,965 of 6,784). Units with intermediate values (0.25–0.3 ms, 228 of 6,784) were excluded. On average, FSs showed narrower waveforms (Extended Data Fig. 2c) and higher firing rate (Extended Data Fig. 2d) than PNs. The classification was verified by delay-epoch photostimulation, during which the mean firing rate of FSs increased due to the activation of PV interneurons, while the mean firing rate of PNs decreased compared to control trials (Extended Data Fig. 2e). Unless stated otherwise, we focused our analyses on PNs with mean firing rate > 1 spike/s.

#### Single neuron activity analysis

Neurons were defined as selective within a specific time window if their spike counts significantly distinguished left and right trials (Mann-Whitney *U* test, *P* < 0.05). An increasing proportion of neurons are selective from the sample, delay to response epochs. In the left ALM of TAs mice (1240 selective neurons), 39.0%, 42.2%, and 58.5% of neurons are selective during the sample, delay, and response epochs, respectively, in the tactile task, while 27.7%, 33.4%, and 51.9% are selective in the auditory task. Bimodal neurons, selective in both tasks, account for 13.4%, 20.5%, and 39.2% across epochs. In the left ALM of TAr mice (1779 selective neurons), 40.3%, 39.0%, and 51.3% are selective across the sample, delay, and response epochs in the tactile task, and 27.4%, 31.9%, and 51.3% in the auditory task; bimodal neurons comprised 13.6%, 16.3%, and 33.2% of neurons. Similar fractions were observed in the right ALM. In each task (tactile or auditory), neurons were classified as left-preferring if their spike counts in left trials were significantly higher than in right trials, or as right-preferring if spike counts in right trials were significantly higher than in left trials. Otherwise, neurons were deemed as non-selective (*P* ≥ 0.05). Furthermore, neurons were classified based on their selectivity in tactile and auditory tasks. Neurons that showed selectivity only in the tactile task were identified as tactile-specific neurons (Extended Data Fig. 3a, d, g, j), while those showing specific selectivity in the auditory task were classified as auditory-specific neurons (Extended Data Fig. 3c, e, h, k). Neurons with selectivity in both tactile and auditory tasks were considered as bimodal neurons (Extended Data Fig. 3c, f, i, l, bimodal neurons showing consistent selectivity in both tasks; See Fig. 2a–b for example neurons of bimodal neurons with contradictory selectivity). In Extended Data Fig. 3a–f, neurons were classified based on their spike counts from sample onset to the end of response epoch. In Fig. 2a–b and Extended Data Fig. 3g–l, neurons were classified with selectivity in delay epoch.

To analyze the dynamic changes in selectivity across neuron types, we selected pyramidal neurons that showed selectivity in at least one epoch of either task, excluding those without selectivity in all epochs of both tactile and auditory tasks. The proportion of each type of selective neurons in a given task epoch was calculated by dividing its count by the total number of selective neurons across all epochs. The changes in selectivity and proportions over epochs were visualized using a Sankey plot (Fig. 2c, d).

To track temporal changes in the proportion of bimodal neurons within a finer time scale (200-ms windows with 20-ms steps), we assessed selectivity in each window and included neurons exhibiting selectivity in at least one window from −0.3 to 3.2 s relative to sample onset. For each window, the proportion was computed as the count of each selective type divided by the total number of selective neurons across all windows (Fig. 2e–g). In Fig. 2e, a black tick was plotted above the middle trial window when TAr showed significantly higher proportions than TAs across three consecutive trial windows. Standard errors were estimated by bootstrapping sessions 1,000 times with replacement, defined as the standard deviation of the proportions across resampled datasets.

To determine the selectivity type of neurons, we fitted single-neuron firing rates (FRs) within specific time windows using three task variables (three-way ANOVA). For sample-epoch and delay-epoch activity, we fitted FRs with task modality (T), sensory instruction (S) and choice (C). For response-epoch activity, we fitted FRs with task modality (T), choice (C) and outcome (O). Trials can be classified into eight conditions based on these variables: tactile left correct trial (TLC), tactile right correct trial (TRC), tactile left error trial (TLE), tactile right error trial (TRE), auditory left correct trial (ALC), auditory right correct trial (ARC), auditory left error trial (ALE) and auditory right error trial (ARE). Only sessions with at least five high-quality PNs recorded simultaneously for at least eight trials of each correct trial type and five trials of each error trial type were included in this analysis.

We fitted a linear regression model of single-trial activity using the MATLAB (2021b) function *fitlm* with the formula:

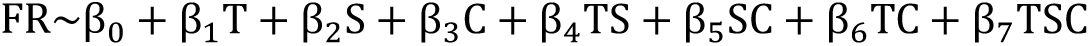

Or

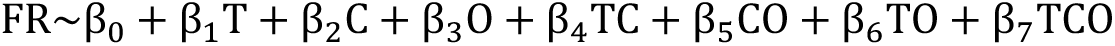

Neurons with significant fitted results (*F*-test, *P* < 0.05) were selected for further classification based on the coefficients (β). Neurons showing significance (*t*-test, *P* < 0.05) in any interaction term (β_4_−β_7_) were classified as nonlinear mixed selective (NMS). Neurons that exhibited significance in exactly one main effect (β1−β3) without significant interaction terms were defined as pure selective (PS). Neurons significant in two or more effects (β_1_−β_3_) but not in interaction terms were classified as linear mixed selective (LMS). Non-selective (NS) neurons showed no significant terms among β_1_−β_7_. The proportions of selective neurons in TAs and TAr were compared using a two-sided permutation test (10^4^ bootstrapping).

#### Population decoding

Decoding of task variables was performed on pseudo-populations collected from different recording sessions. Only sessions with at least eight trials of each trial type (both correct and error) were used for this analysis. Trials during task transitions were excluded from this analysis to prevent contamination from the unstable neural coding ability in these trials.

For each time window, linear support vector machines (SVMs) were independently trained using single-trial activity to decode sensory instruction, choice, outcome, and task modality (MATLAB function *fitcsvm*). The number of trials was matched across all trial types. Within-condition decoding accuracy was estimated by either 10-fold cross-validation (Fig. 3, Extended Data Fig. 4) or prediction on a test set that matched the training set in trial number (Extended Data Fig. 5), with both approaches yielding consistent results. Cross-condition decoding accuracy was assessed by measuring classification accuracy on the other condition (as indicated by the dashed arrows in Extended Data Fig. 4c). Decoding was repeated 100 times, with random sampling of neurons and trials in each run. Decoding performance was presented as mean ± s.d. across runs. Chance levels were estimated by shuffling training set labels to generate a null model.

To estimate the decoding performance with increasing neuron numbers (Extended Data Fig. 4a–b), each decoder was trained and tested on 200-ms time-averaged single-trial neural activity. Time windows for sensory instruction, choice, outcome, and task modality were set to 0 to 0.2 s (early sample epoch), 1.8 to 2 s (late delay epoch), 2.4 to 2.6 s (mid response epoch) and −0.2 to 0 s (pre-sample period), respectively. In other figures, the pseudo-populations were constructed from 300 randomly chosen neurons.

To perform cross-temporal decoding (Extended Data Fig. 5e–h), decoders were trained on 200-ms time-averaged neural activity ranging from −1 to 4 s with 20-ms steps, and then tested on activity across all time windows. For the test window that matched the training window, independent trials were used for training and testing. Decoding was repeated 10 times and averaged across runs. The results showed an abrupt change in the representational subspace following go cue, revealing that distinct neural populations encoded sensory-related information in the sample and response epochs, and choice-related information in the delay and response epochs. Moreover, the peak of “sensory” decoding accuracy occurred later than both the lick latency and outcome decoding (Extended Data Fig. 5i, j), reflecting a post-outcome feedback of sensory instruction.

Cross-condition generalization performance (CCGP) quantifies the generalizability of neural representations across contexts^45,54^. As illustrated in Extended Data Fig. 4c, training and testing sets were swapped to generate two values of cross-condition decoding accuracies, with the overall CCGP computed as their mean. Similarly, the general within-condition decoding performance (WCDP) was calculated as the average within-condition decoding accuracy across datasets. To quantify the CCGP relative to WCDP, a generalization index (GI) was computed as follows^55^:

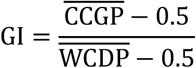

In this formula, 0.5 represents chance level. CCGP and WCDP refer to the average CCGP and WCDP within specific time windows, respectively. The time window for sensory instruction GI is 0 to 1 s, for choice GI is 1 to 2 s, for action GI is 2 to 2.3 s, for outcome GI is 2.2 to 3 s, and for task modality GI is −1 to −0.5 s (Fig. 3c). A GI closer to 1 indicates that the neural representations can be generalized across conditions at a level comparable to within-condition performance, whereas a GI closer to 0 indicates generalizability approaches chance. GIs of TAs and TAr were compared using a bootstrap test for equality of means (repeated 1,000 times to generate the null distribution^55^). .

#### Decomposition of population activity using targeted dimensionality reduction

Neuronal population activity can be decomposed into several activity modes to capture distinct features^19,39^. Same as decoding, only sessions with at least eight trials of each trial type (both correct and error) were included. To retain neurons with stable firing during task, we computed the Pearson correlation coefficient of neuronal activity between the two halves of correct trials for individual neurons. Neurons with a correlation coefficient of at least 0.5 were selected for further analysis.

Activity modes were calculated using 75% randomly chosen trials, leaving the remaining trials for projections. Trials during task transitions were excluded. Specifically, for each trial type, we calculated time-averaged and trial-averaged population activity vector (𝑛 × 1, where *n* is the number of neurons), named as ***TLC***, ***TRC***, ***TLE***, ***TRE***, ***ALC***, ***ARC***, ***ALE***, ***ARE***. Modes for tactile and auditory tasks were separately calculated.

To obtain sensory mode that captures the saliency of sensory inputs, we averaged neural activity during sample epoch (0 to 1 s). The tactile sensory modes for TAs and TAr were calculated as:

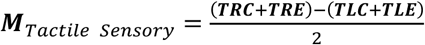

Due to the reversed sensory-motor contingency in auditory task of TAr, the auditory sensory modes for TAs and TAr were calculated in opposite ways:

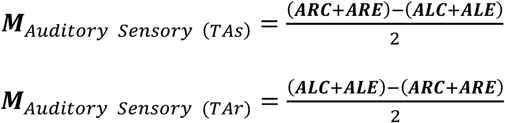

For other activity modes, tactile and auditory tasks followed the same calculations, we used ***LC***, ***RC***, ***LE*** and ***RE*** to refer to task-general activity vectors. We used delay epoch-averaged activity (1 to 2 s) to calculate choice mode and early response epoch-averaged activity (2 to 2.3 s) to calculate action mode:

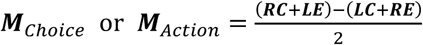

Outcome mode was calculated using response epoch-averaged activity (2 to 3 s):

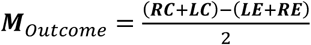

In addition to these selective modes that can distinguish trial types based on differential task variables, there were activity modes capturing non-selective activity features. The ramping mode was defined by the difference between the average activity during late delay period (1 to 1.5 s) and that during pre-sample period (−0.5 to 0 s). It captured the non-selective ramping activity over time. The go mode was calculated as the difference in activity between the time after go cue (2 to 2.1 s) and before go cue (1.9 to 2 s).

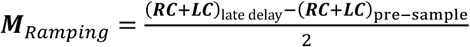

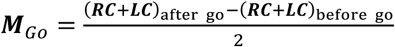

Each activity mode (***Mode***, 𝑛 × 1) was normalized by its norm. Population activity projection along each activity mode were calculated as ***Mode**^T^**r***, where ***r*** is the trial-averaged population activity matrix (𝑛 × 𝑡, where *t* represents the number of time points in a single trial). This calculation was repeated 10 times, with trials randomly sampled for activity modes and projections in each run. Projection was presented as mean ± s.d. across runs.

To estimate the relationships among activity modes in the activity space, we evaluated the cosine of angle between each pair of modes (cos θ). A value closer to 0 indicates that the two directions are nearly orthogonal to each other, suggesting that they tend to use different groups of neurons. The values in Fig. 3d–e were the averages across 10 runs. To assess whether tactile and auditory modes are more parallel in the TAs than in the TAr contingency, we compared the observed difference (cos θ in TAs − cos θ in TAr) with a null distribution, which is generated by shuffling the neuronal weights on each activity mode within task and within contingency and then recalculated cos θ between the random activity modes over 10^4^ repetitions. The *P* value was defined as the probability that the null distribution was higher than the observed difference (one-sided test).

#### Comparison of neuron weights on activity modes

Each activity mode represents a vector of neuron weights. To quantify and compare neuron weights on tactile and auditory activity modes, we calculated the mean activity mode for each mode by averaging over runs, followed by re-normalization. This yielded a vector of mean neuron weights, representing each neuron’s contribution to the corresponding mode.

For each specific task component (***M****_Intensity_*, ***M****_Choice_*, ***M****_Action_*, ***M****_Outcome_*, ***M****_Ramping_* and ***M****_Go_*), we calculated Pearson correlation coefficients (ρ) between weights on the tactile mode (W_Tac_) and the auditory mode (W_Aud_). To further compare coefficients in TAs (ρ_s_) and TAr (ρ_r_), we calculated their difference as ρ_s_−ρ_r_, and compared it with a random null distribution. Specifically, neuronal weights on each activity mode were shuffled within task and within contingency over 10^4^ repetitions. We then calculated the random distribution of differences ρ_s_,_random_−ρ_r_,_random_. The *P* value was defined as the probability that the random distribution was higher than ρ_s_−ρ_r_ (one-sided test). We also tried an alternative shuffling method, where we first generated a random dataset of W_Tac_ and W_Aud_ independently. Taken W_Tac_ as an example, let n_s_ and n_r_ represent the neuron counts in TAs and TAr datasets, respectively. We randomly chose n_s_ samples from the overall (n_s_+n_r_) samples as the random weights in TAs, leaving the remaining n_r_ samples as the random weights in TAr. Similarly, we generated the random weights for W_Aud_. We then calculated ρ between the random tactile and auditory weights, i.e. ρ_s_,_random_ and ρ_r_,_random_ for TAs and TAr datasets, respectively. This process was repeated 10^4^ times with bootstrapping to obtain the random distribution of differences ρ_s_,_random_−ρ_r_,_random_. In each repeat, the dataset of (n_s_+n_r_) samples was generated by randomly sampling with replacement from neuron weights in TAs and TAr. This approach yielded similar *P* values—0.0396, 0.0016, 0.3587, 0.1486, 0.0042 and 0.1786 for ***M****_Intensity_*, ***M****_Choice_*, ***M****_Action_*, ***M****_Outcome_*, ***M****_Ramping_* and ***M****_Go_*, respectively—comparable to those from the first approach: 0.0336, 0.002, 0.3629, 0.1561, 0.0063 and 0.1683.

Neuron weights on each activity mode were visualized in the *t*-SNE presentation following ref. ^19^. Tactile and auditory activity modes were concatenated as an 𝑛 × 12 matrix for *t*-SNE. We ran *t*-SNE 10 times, with the perplexity of 50 and correlation distance. The two-dimensional embeddings with the lowest Kullback-Leibler divergence were used for visualization.

#### ePAIRS test

We tested whether neuronal activity patterns are uniformly distributed over the representational space or clustered as functional population modules, using both the response matrix and the activity mode matrix. The response matrix was constructed as an 𝑛 × 𝑝 matrix, where the rows represented the baseline-subtracted, 200-ms time-smoothed and magnitude-normalized PSTHs of individual neurons, with TLC, TRC, ALC, ARC trials concatenated. We reduced the dimensionality using PCA, retaining 99% of the variance. For a de-noised matrix that is 𝑛 × 𝑑, each neuron’s activity over time is represented as a 1 × 𝑑 vector. We calculated the average angle between the activity vector of each neuron and its k nearest neighbors, and then obtained the distribution of average angles across all n neurons. To generate the null distributions that captured the variance structure of neural data, we followed ref.^18,19^. The distributions were constructed as 𝑛 × 𝑑, with each neuron’s activity was a d-dimensional Gaussian. We calculated the null distributions 10^4^ times and compared them with the empirical distribution using Wilcoxon rank sum test. For the activity mode matrix, which is an 𝑛 × 12 matrix, each neuron was featured as a 12-element vector without dimensionality reduction. The ePAIRS test employed the same procedure as described above (results shown in Extended Data Fig. 7a, Extended Data Fig. 8a). Both approaches consistently reveal significant functional clustering.

#### Multidimensional scaling

We applied multidimensional scaling (MDS) to map the neuronal population activity into a 3D space. This method preserves the spatial relationships (typically distances) among task conditions. For each task epoch (sample, delay, and response epochs), we obtained time-averaged and trial-averaged activity for each trial type (TLC, TRC, TLE, TRE, ALC, ARC, ALE, ARE) and then constructed a response matrix of size 8 × 𝑛, with each column z-score normalized. We calculated the correlation distance between these trial types. The correlation distance was then subjected to non-classical MDS using the MATLAB function *mdscale*, with the criterion set to ’metricstress’.

#### Quantifications of representational dimensionality

We estimated the dimensionality using neural activities in correct trials. Sessions with fewer than 20 correct trials were excluded for this analysis. The method for quantifying dimensionality followed ref.^5^, which is based on the observation that the number of implementable binary linear classifiers grows exponentially with the increasing number of dimensions. For a specific number of neurons, we estimated the maximal number of conditions for which 95% of all the binary classifiers are implemented. The implementable classifiers were those showing at least 75% cross-validated classification accuracy. The maximal dimensionality is bounded by the number of trial conditions (four trial types: TLC, TRC, ALC and ARC).

#### Roles of different neurons on dimensionality

To validate roles of mixed selectivity on representational dimensionality, we removed different neuron types from the neural population and estimated dimensionality of the remaining neurons. After excluding one (or two) specific neuron types, the remaining neurons were randomly sampled while preserving their relative proportions within the new group in each iteration. For fair comparison, the dimensionality of the original full neuronal population was also re-estimated with neuron type proportions preserved in each iteration. Dimensionality was assessed as a function of the number of neurons as described above (see Quantifications of representational dimensionality). The relative effect of neuron type exclusion was quantified by the relative area under the curve (relative AUC), calculated by subtracting the mean AUC of the full population from the AUC of each trace (each iteration). Since removing different neuron types results in varying population sizes, we calculated AUCs in each panel using the minimum population size across all conditions as the upper bound (Fig. 4g, h, Extended Data Fig. 10d, e).

To compare the dimensionality after controlling for differences in neuron type composition between contingencies, we matched the proportions of NMS, LMS, PS, and NS neurons in the TAr dataset to those in the TAs dataset (Fig. 4i, Extended Data Fig. 10f). To further rule out the effect of differences in specific NMS subtypes, we additionally matched the composition of NMS subcategories that differed between contingencies across datasets. These finer-level adjustments did not significantly affect the results.

#### Decomposition of population activity during task transition

To decompose the neural representations during task transition, we projected single-trial population activity during transition along each activity mode for each session. For the single-session analysis of task transition, we applied a more lenient criterion to select sessions. Specifically, we included sessions with at least five high-quality PNs recorded simultaneously, with a minimum of eight trials for each correct trial type and six trials for each error trial type (Some sessions did not have enough error trials, especially in TAs mice).

For each session, we constructed population activity matrices for left and right trials independently during the task transition, excluding optogenetic trials, pre-licking trials and no-response trials. The neural activity was averaged within specific time windows: 0 to 1 s for ***M****_Intensity_*, 1 to 2 s for ***M****_Choice_*, 2 to 2.3 s for ***M****_Action_*, and 2 to 3 s for ***M****_Outcome_*. For each activity mode, the activity matrices of transition trials for projection were denoted as ***V_L_*** and ***V_R_*** for left and right trials, with sizes of 𝑛 × 𝑞_1_ and 𝑛 × 𝑞_2_, where *n* is the number of neurons, and *q_1_* and *q_2_* are the numbers of left and right trials, respectively.

Given the smaller number of neurons for single-session analysis, we randomly sampled 50% of the trials as the training set, leaving the other half of trials for the test set to ensure stable test scores. We computed the tactile and auditory activity modes as described above (see Decomposition of population activity, but here for individual sessions). Furthermore, we applied the Gram–Schmidt process to these activity modes to ensure they were fully orthogonal to one another. Each mode was then normalized by its magnitude. The single-trial projections were calculated as ***Mode***^*T*^***V_L_*** and ***Mode***^*T*^***V_R_***, followed by baseline subtraction, where the baseline was defined as the projections of pre-sample activity from −1 to −0.5 s. Test scores, which reflected the projections of stable trials, were derived from the projections of trial-averaged activity in the test set and then baseline-subtracted. This procedure was repeated 20 times, and the results were averaged to obtain the mean projections for each session.

Sessions were categorized into two groups based on their transition types: tactile-to-auditory and auditory-to-tactile transitions. Projection data from the same transition type was grouped for visualization. Although the number of trials required for successful transitions varied across sessions, the overall trends in behavioral changes were similar. The single-trial activity projections from different sessions were aligned to the first transition trial (with left and right trials arranged separately). We used a 15-trial window with 1-trial step to calculate the hit rate (mice’s performance) and to smooth the projections over trials.

#### Decoding choice during task transition

To estimate the choice information decodability by neuronal populations during task transition (Fig. 6g–i, Extended Fig. 11g–j), we trained the linear SVMs using stable tactile and auditory trials as described above (see Population decoding but for individual sessions), and tested on single-trial activity throughout transition. Sessions with at least five high-quality PNs recorded simultaneously and at least six trials of each trial type (both correct and error) were included for this analysis. For each session, the population activity matrices were constructed as described above (see Decomposition of population activity during task transition). The time window for decoding choice was 1 to 2 s relative to sample onset.

For each session, decoders were trained with 100 repeats, with six trials randomly sampled for each trial type in each repeat. We then tested their decoding performance on the single-trial activity matrices with a 15-trial sliding window and a 1-trial stride. In Fig. 6g–i, we applied choice decoders to classify the sensory instruction of each trial, assessing how effectively the neural population could support task execution. In Extended Fig. 11g–j, we evaluated their performance on classifying choice over trials, to test whether the choice representation was fully engaged. The final result was obtained by averaging performance across repeats.

Sessions were categorized into two groups following the aforementioned procedures. Decoders trained on left and right ALM neurons exhibited similar decoding performance (not shown here). Therefore, in the presented results, we trained the decoders on combined data from both hemispheres to enhance decoding stability.

For behavioral and all the decoding performance, we defined recovery window as the first trial window post-transition where the left and right performance reached an equal level (i.e., converged).

To enable comparison of behavioral and neural recovery after transition (Fig. 6g–i), behavioral and neural performance were calculated using the same window size (15-trial) and step size (1-trial). To estimate the variance of transition recovery, we conducted a leave-one-session-out analysis, in which each run excluded one session and recalculated the behavioral and neural recovery windows (Fig. 6i).

To evaluate the overall trend of choice decoder performance during task transition and compare the decoding accuracy between left and right transition trials (Extended Fig. 11g–j), we estimated the average trend of decoding accuracy change. Specifically, we first calculated the secant slope, defined as the slope of the line connecting each trial window to the first transition trial window. The average trend of decoding performance change was then quantified as the mean secant slope over all trial windows spanning from the transition window to the recovery window.

## Extended Data Figures

**Extended Data Fig. 1.**
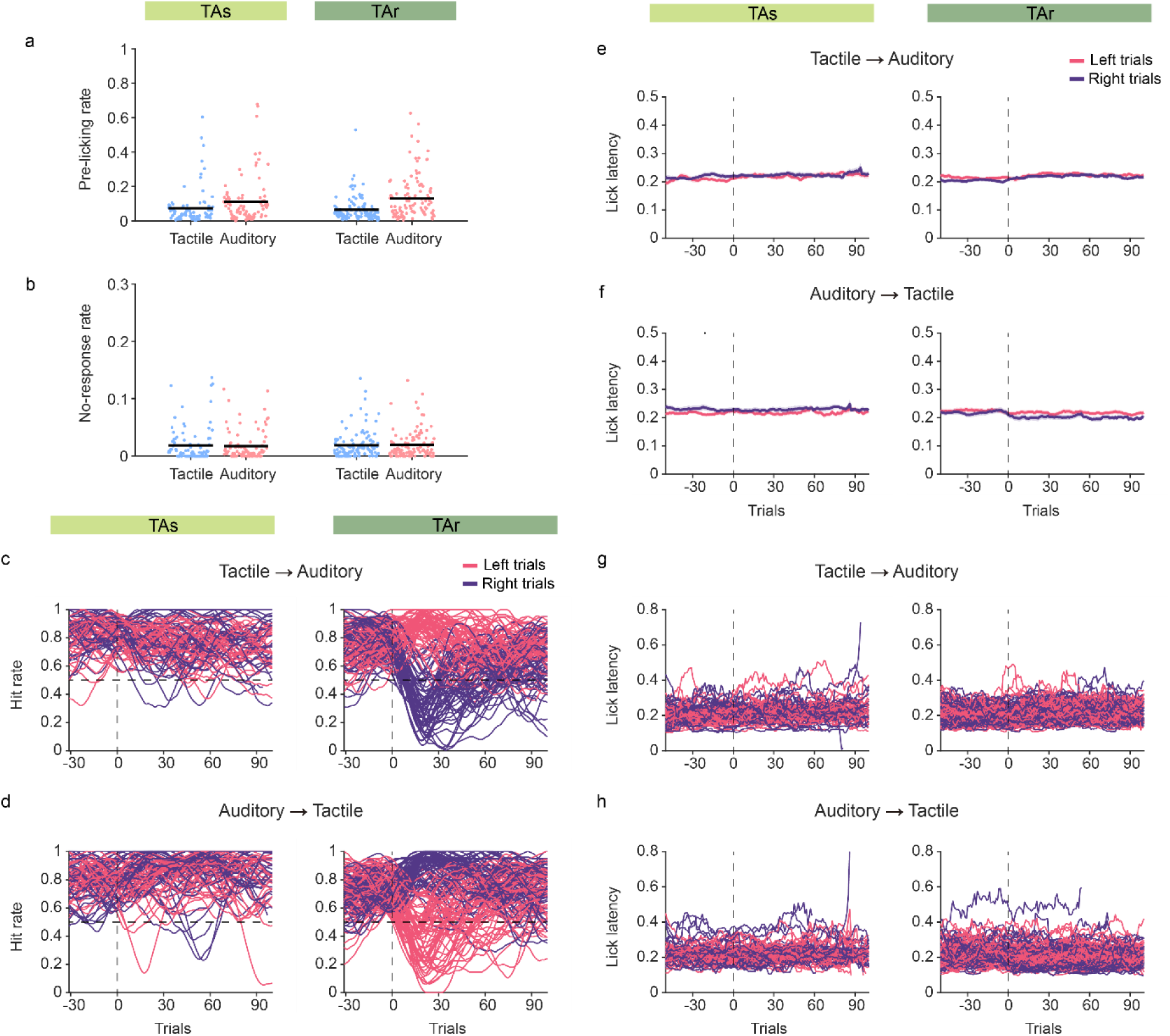
Behavioral analysis for expert mice. **a**, Pre-licking rate of stable trials. Dots, single sessions; black bar, mean. **b**, No-response rate of stable trials. **c, d**, Same as Fig. 1 c, d, but for visualization of single sessions. **e, f**, Lick latency during tactile-to-auditory transition and auditory-to-tactile transition. No significant differences between latencies of left and right trials were found during task transition. Mean ± s.e.m. (bootrstrap) across sessions. **g, h**, Same as e, f, but for visualization of single sessions.

**Extended Data Fig. 2.**
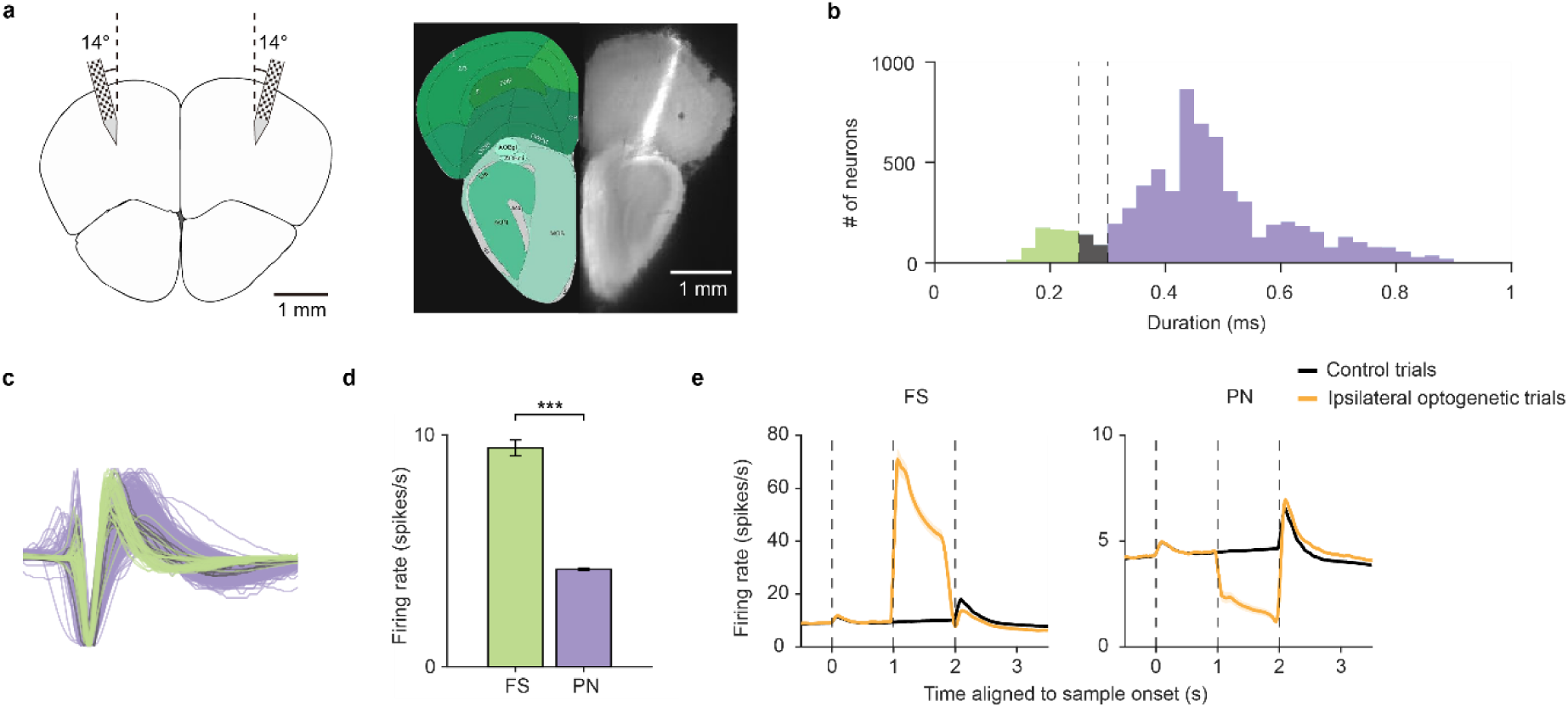
Electrophysiological recording with Neuropixels probes and classification of pyramidal and fast-spiking neurons. **a**, Simultaneous recording of the left and right ALM using Neuropixels 1.0 probes. Left, schematic of probe insertion. Right, example recording session showing probe trace labeled by CM-DiI. **b**, Distribution of spike durations, measured by trough-to-peak width. Only neurons with firing rate > 1 spike/s and duration < 0.9 ms were included. Putative fast-spiking neurons (FS, green, n = 591) and putative pyramidal neurons (PN, purple, n = 5,965) were classified based on their durations. 228 neurons were not classified (gray). **c**, Spike waveforms of 1,000 randomly sampled neurons with the same proportions as in b. **d**, Averaged firing rate of classified FSs and PNs, mean ± s.e.m. (bootstrap). \*\*\**P*<0.001, two-sample t-test. **e**, Averaged PSTHs of FSs and PNs in all control trials and delay-epoch ipsilateral optogenetic trials.

**Extended Data Fig. 3.**
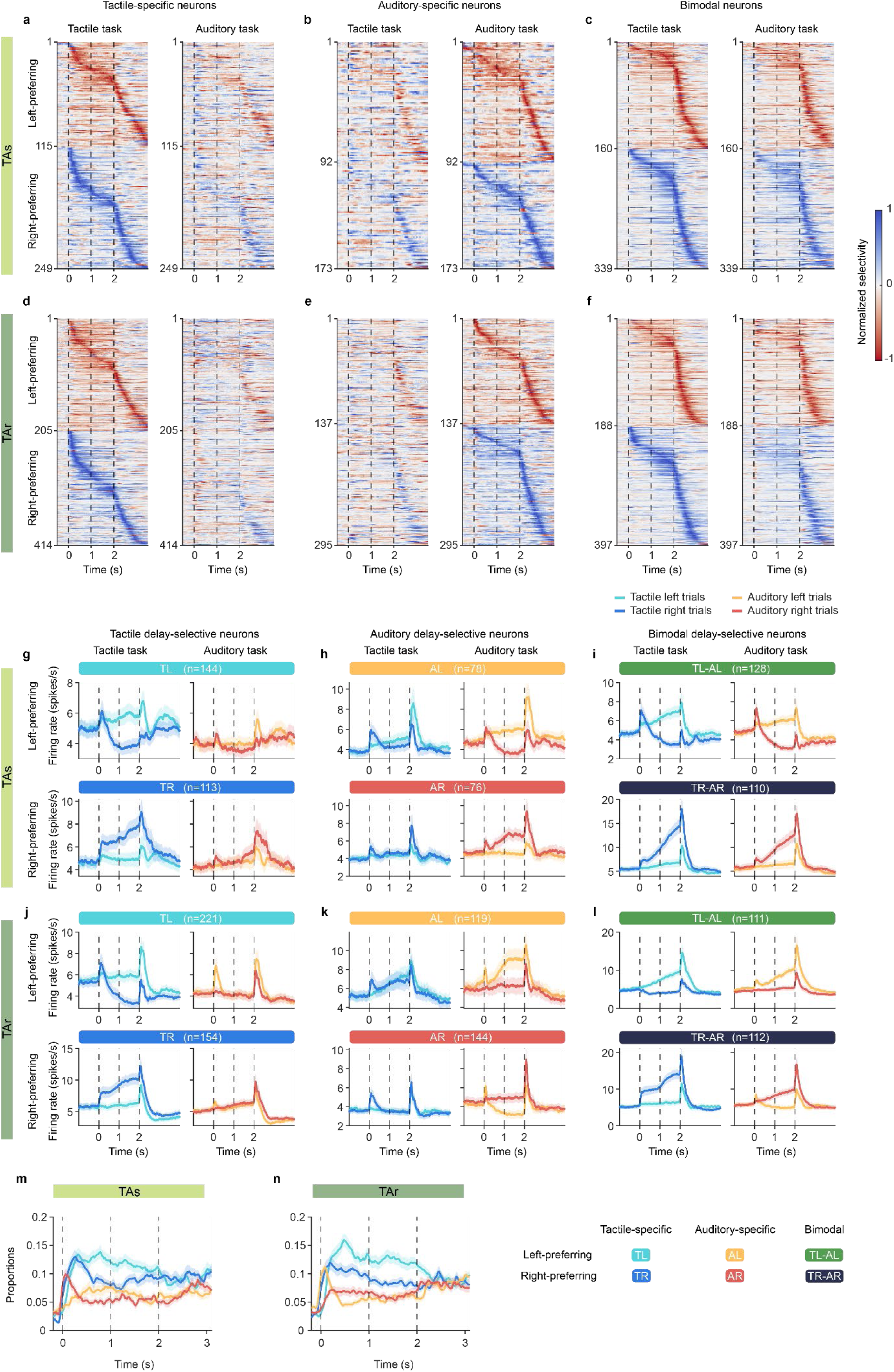
Classification of selective neurons. **a**, Normalized selectivity of tactile-specific neurons of TAs mice. These neurons showed selectivity in tactile task (left column) but not in auditory task (right column). Left-preferring (red) and right-preferring (blue) neurons were sorted by their peak selectivity in tactile task. **b**, Same as a, but for auditory-specific neurons. Neurons were sorted by their peak selectivity in auditory task. **c**, Same as a, but for bimodal neurons. Neurons were sorted by their peak selectivity in tactile task, and they also showed selectivity in auditory task. **d–f**, Same as a–c, but for neurons of TAr mice. **g**, Average PSTHs of tactile specific neurons of TAs mice; Upper panel, tactile-specific left-preferring (TL) neurons; Lower panel, tactile-specific right-preferring (TR) neurons. **h**, Same as g, but for auditory-specific neurons. **i**, Same as g, but for bimodal neurons. **j–l**, Same as g–i, but for delay-selective neurons of TAr mice. In a–f, neurons were classified based on their selectivity over all epochs (0–3 s relative to sample onset). In g–l, neurons were classified by their delay-epoch (1–2 s relative to sample onset) selectivity. m–n, Same as Fig. 2f–g, but for proportions of each type of task-specific neurons. All the neurons were recorded from the left ALM.

**Extended Data Fig. 4.**
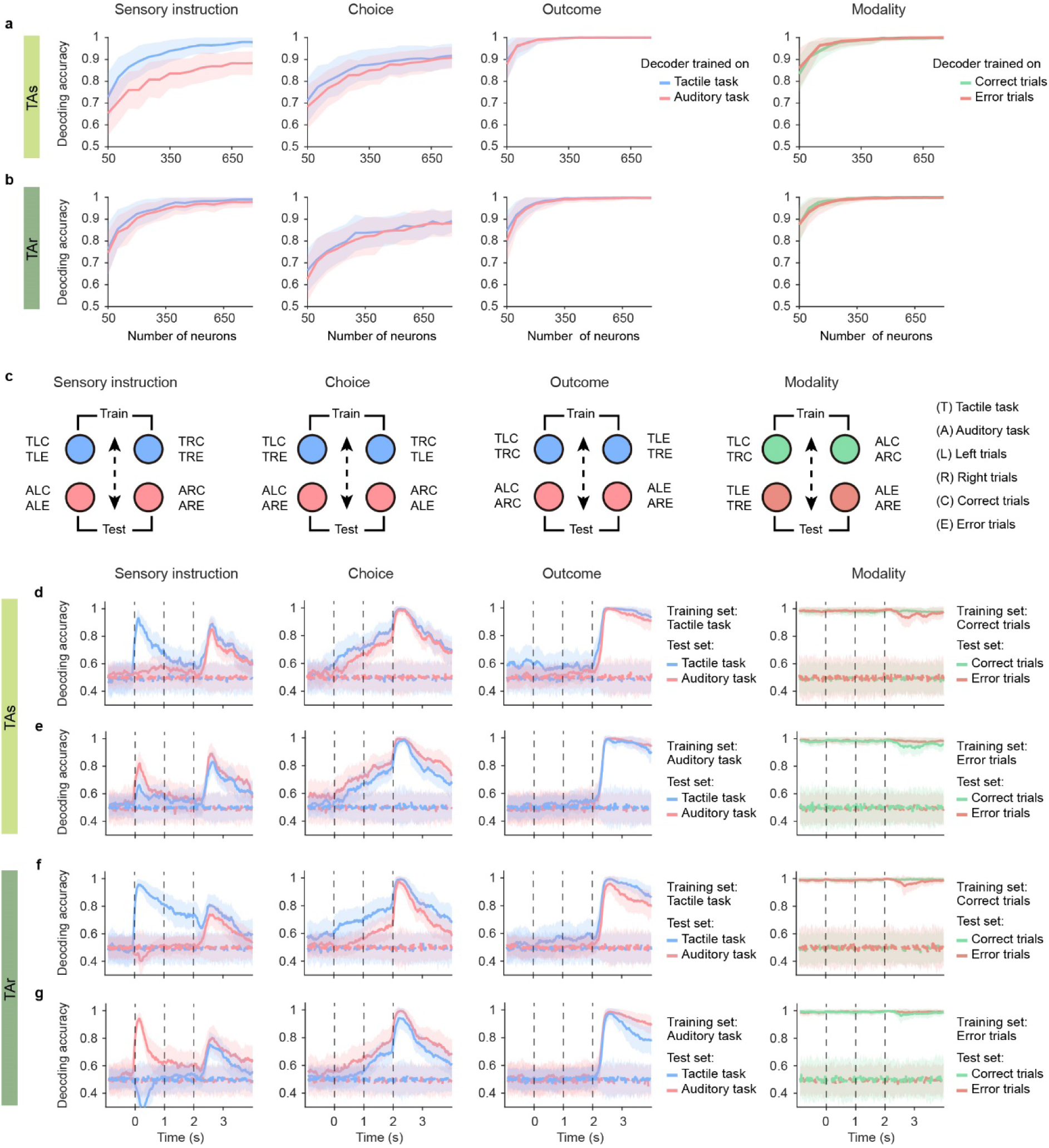
Within-condition and cross-condition decoding analysis. **a**, Decoding accuracy as the number of neurons increased in TAs mice. The accuracy gradually improved with more neurons in the pseudo-population. **b**, Same as a, but for TAr mice. **c**, Schematics that show how linear decoders classify trials into dichotomies and how cross-condition decoding are performed. Dashed arrows separate trials with different labels. Taking the sensory decoder as an example, both LC and LE correspond to left instruction, while both RC and RE correspond to right instruction. Different colors indicate different datasets (conditions). For cross-condition decoding, decoders were trained on one condition and tested on the other condition, and vice versa. **d**, Sensory, choice and outcome decoders trained on tactile tasks were tested on tactile (blue) and auditory (pink) tasks. Modality decoder trained on correct trials was tested on correct (green) and error (orange) trials. Solid line, decoder trained on true data; dashed line, decoder trained on label-shuffled data. **e**, Same as d, but for sensory, choice and outcome decoders trained on auditory tasks, and modality decoder trained on error trials. d–e, for TAs mice. **f**–**g**, same as d–e, but for TAr mice.

**Extended Data Fig. 5.**
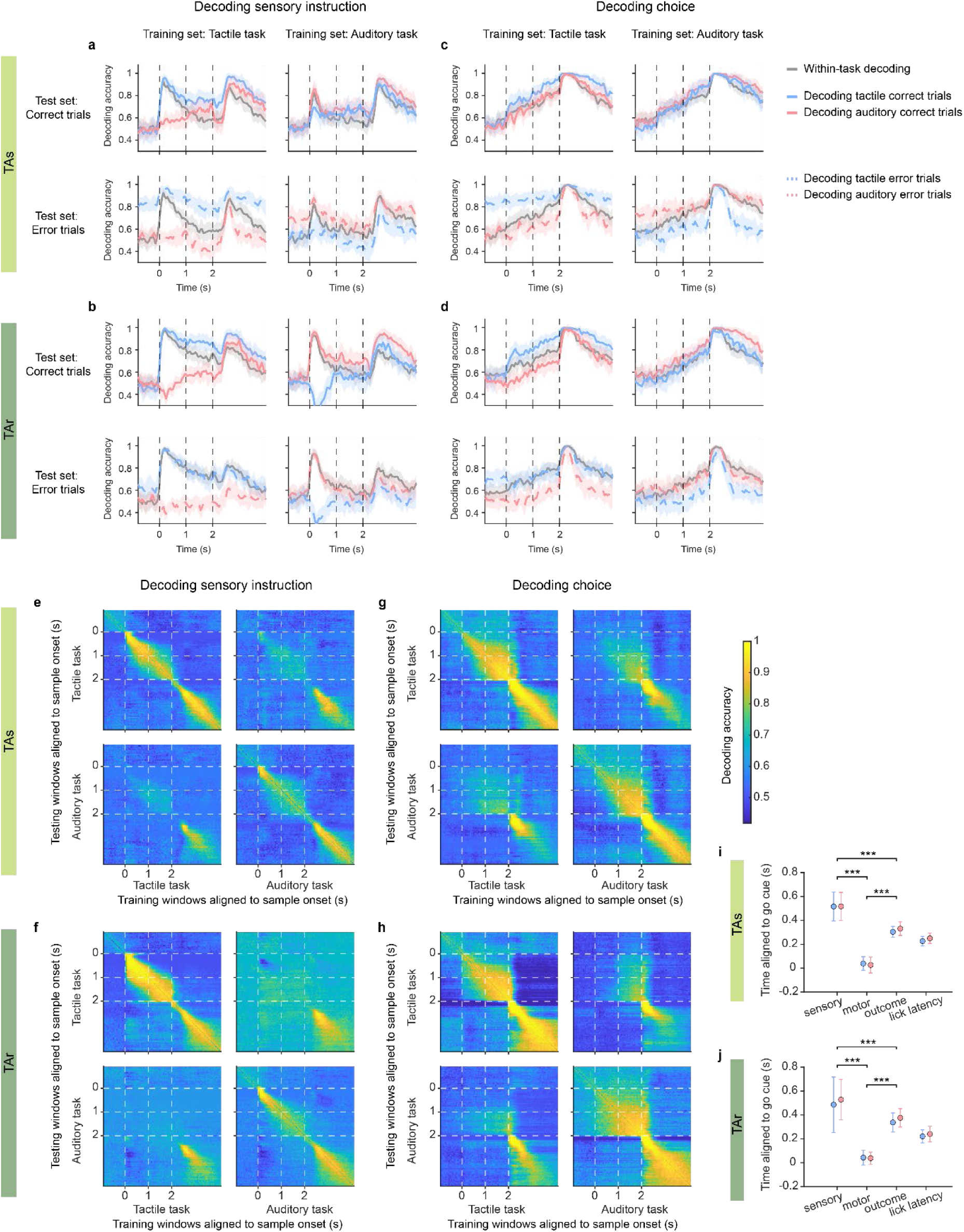
correct-and error-trial decoding and cross-temporal decoding performance. **a**–**d**, Decoding performance of sensory and choice decoders on correct and error trials. **a**, Performance of sensory decoder in TAs mice. Left column, decoder trained on tactile task; right column, decoder trained on auditory task. Top, performance on correct trials (solid line); bottom, performance on error trials (dashed line). Gray, within-task decoding performance. **b**, Same as a, but for TAr mice. **c**–**d**, Same as a–b, but for choice decoders. Decoders showed comparable peak accuracy in correct and error trial during the response epoch. **e**–**h**, Cross-temporal decoding performance of sensory and choice decoders. **e**, Performance of sensory decoder in TAs mice. Upper left and lower right panels, within-task cross-temporal decoding; Note that sample-epoch sensory decoders could not decode sensory information in the response epoch, and vice versa. Lower left and upper right panels, cross-task cross-temporal decoding. Only the sensory instruction encoded during the response epoch could be generalized across tasks. **f**, Same as a, but for TAr mice. **g**–**h**, Same as e–f, but for choice decoders. Note that the abrupt change after the go cue, indicating an intrinsic change in the neural representation of choice following movement initiation. **i** – **j**, Quantification and comparison of the peak times of decoding accuracy and the lick latency. During the response epoch, the choice (motor) decoder reached peak performance first, followed by the mice’s licking behavior. Subsequently, neural activity reflected the outcome of animals’ choices (outcome decoder), and finally, the peak representation of sensory instruction (sensory decoder) emerged. This temporal sequence suggests that sensory information during the response epoch was related to recall or correction of this trial following outcome feedback. \*\*\**P*<0.001, bootstrap test for equality of means.

**Extended Data Fig. 6.**
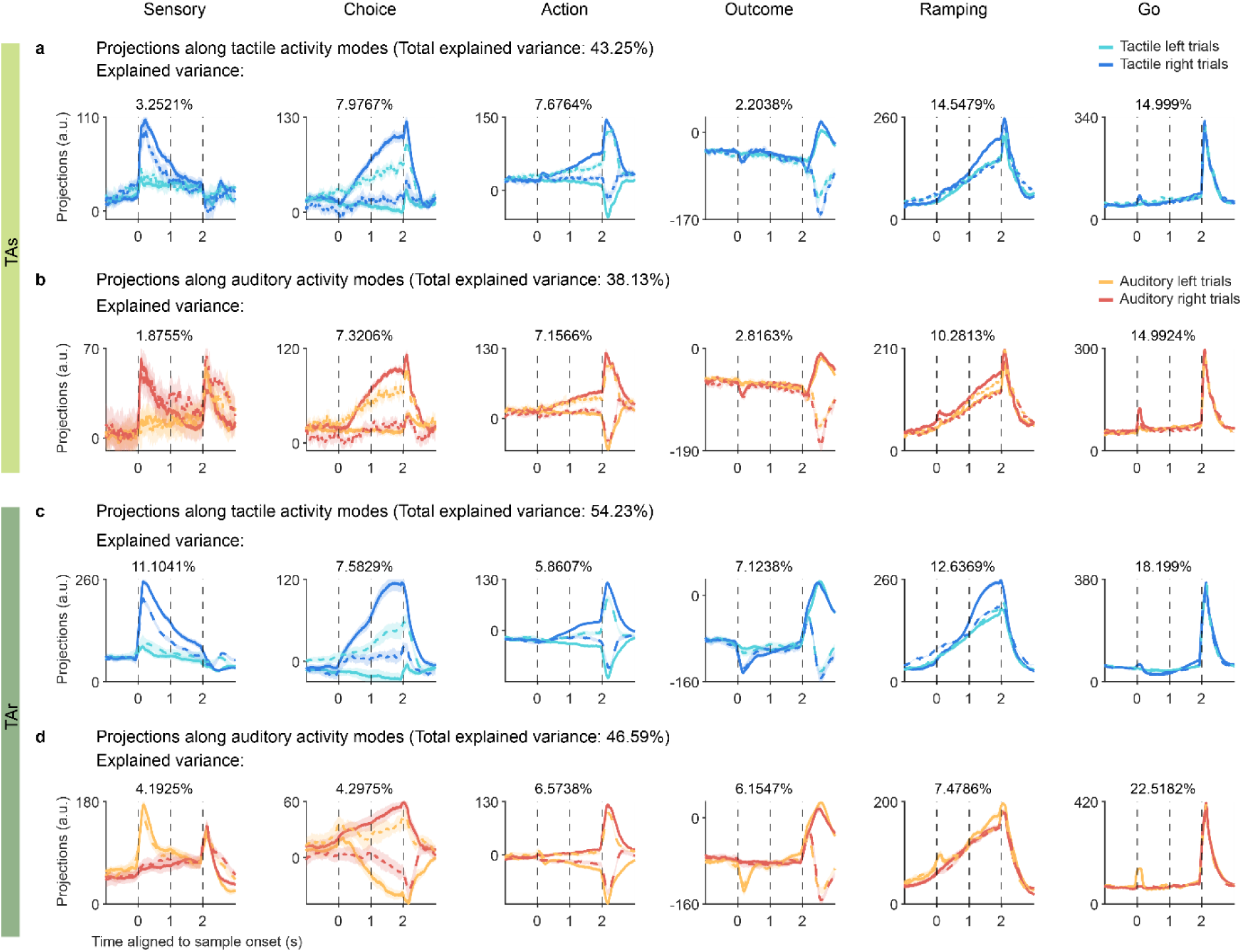
Projections of population neuronal activity along activity modes. **a**, Projections of population activity in tactile trials along tactile activity modes in TAs mice. Solid line, projections of correct-trial activity; dashed line, projections of error-trial activity. Activities along the sensory mode discriminate tactile stimuli, with correct and error trials exhibiting similar projections. Activities along the choice mode and action mode differentiate leftward and rightward choices and movements during the delay and response epochs, respectively; for instance, both the correct right trial and error left trials show projections representing rightward choice. Outcome mode separates correct and error trials, without distinguishing between the left and right trials. Ramping mode captures the increase in activity during the delay epoch relative to the pre-sample epoch, while the go mode signifies the abrupt rise in activity following the go cue. Variance explained by each activity mode is labeled at the top; total variance explained was calculated after mode orthogonalization. **b**, Same as a, but for projections of population activity in auditory trials along auditory activity modes. **c–d**, Same as a–b, but for TAr mice.

**Extended Data Fig. 7.**
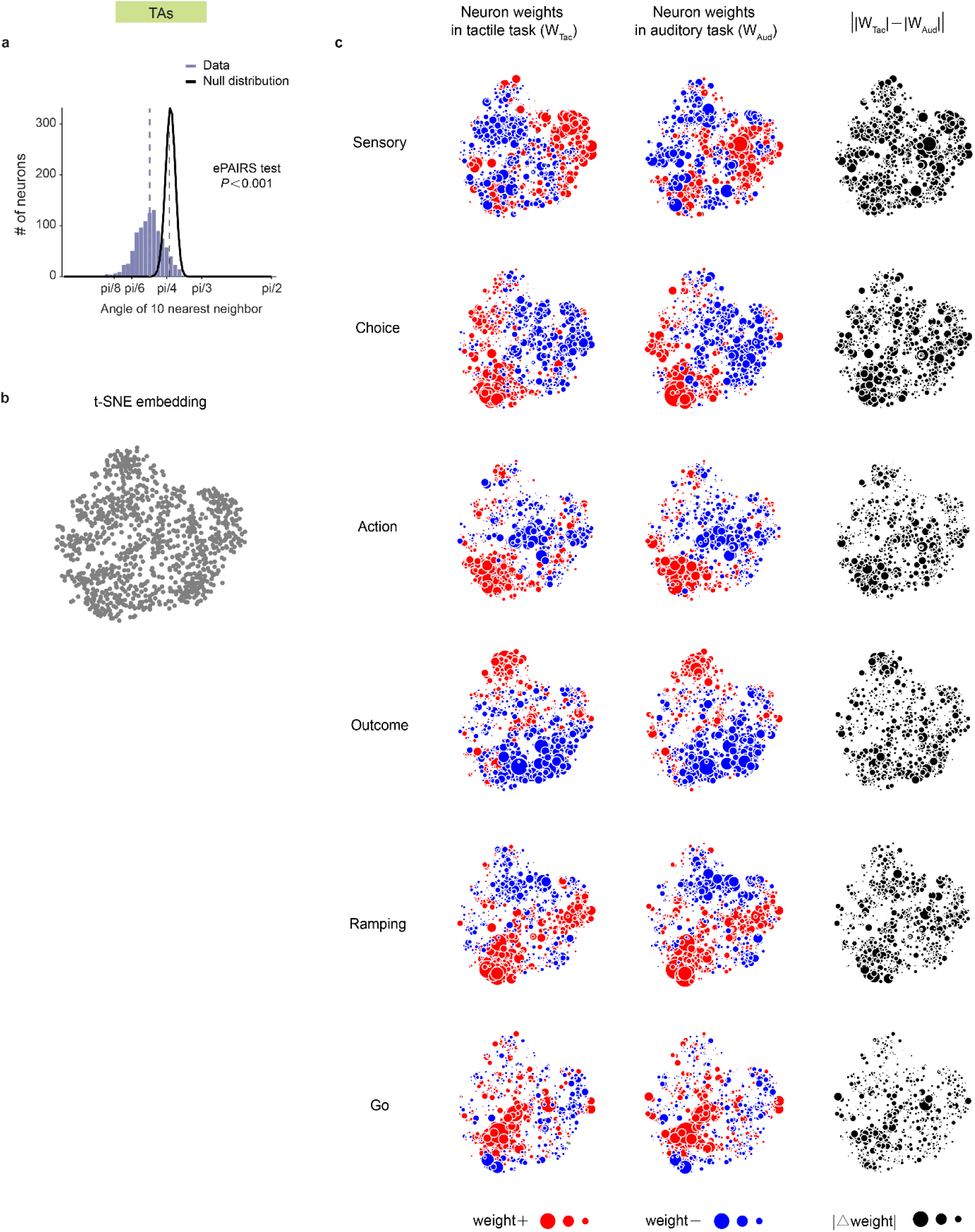
t-SNE visualization of neuron weights for TAs mice. **a**, ePAIRS test, *P* value, two-sided Wilcoxon rank sum test. **b**, t-SNE embedding of all left ALM neurons. **c**, Visualization of neuron weights. Left column, weights on tactile modes (W_Tac_); middle column, weights on auditory modes (W_Aud_); right column, absolute difference between weights on different tasks (||W_Tac_|−|W_Aud_||). The size of each scatter depicts the magnitude of the neuron’s absolute weight. Red, positive weights; blue, negative weights.

**Extended Data Fig. 8.**
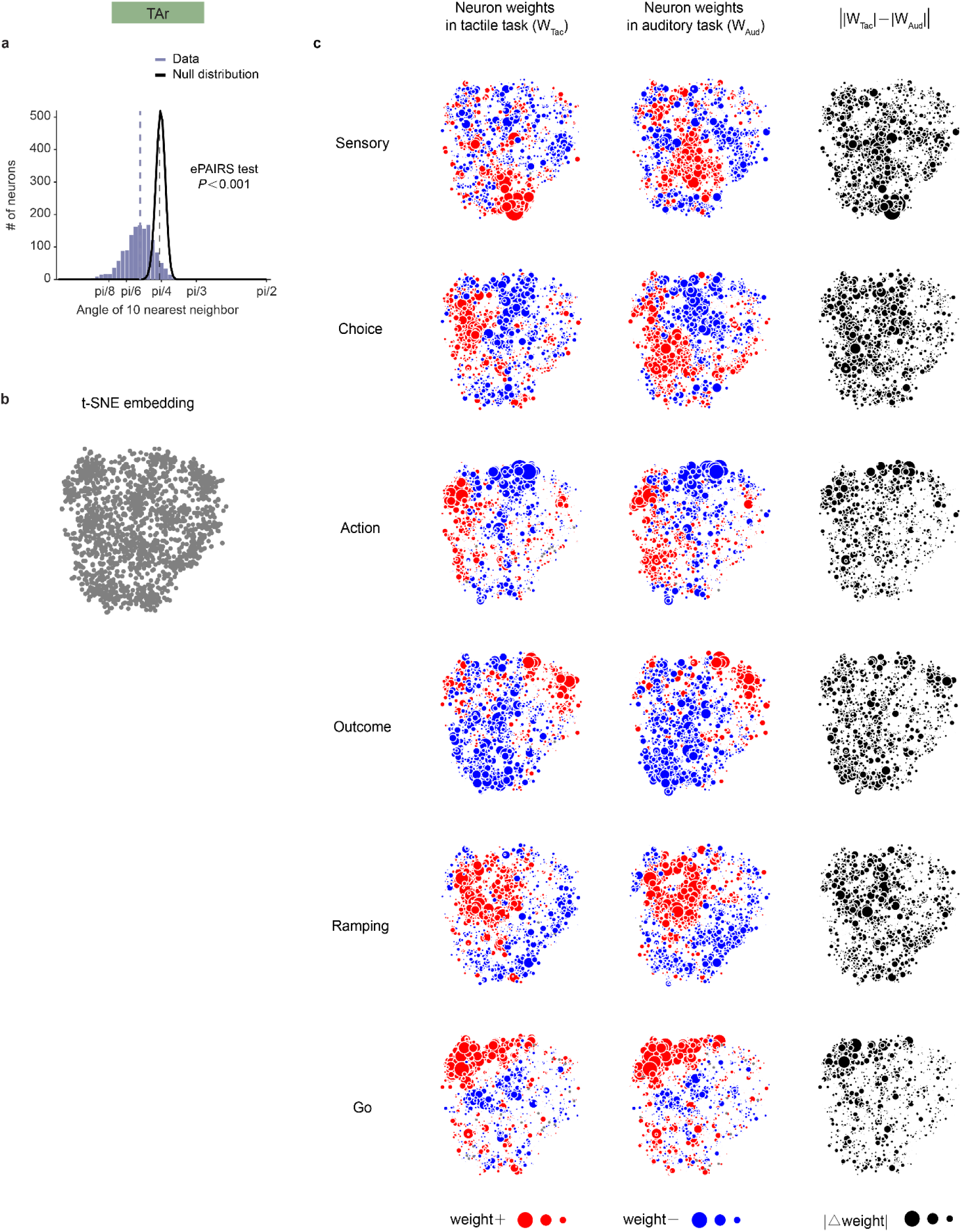
t-SNE visualization of neuron weights for TAr mice. Same as Extended Data Fig. 7, but for the left ALM neurons in TAr mice.

**Extended Data Fig. 9.**
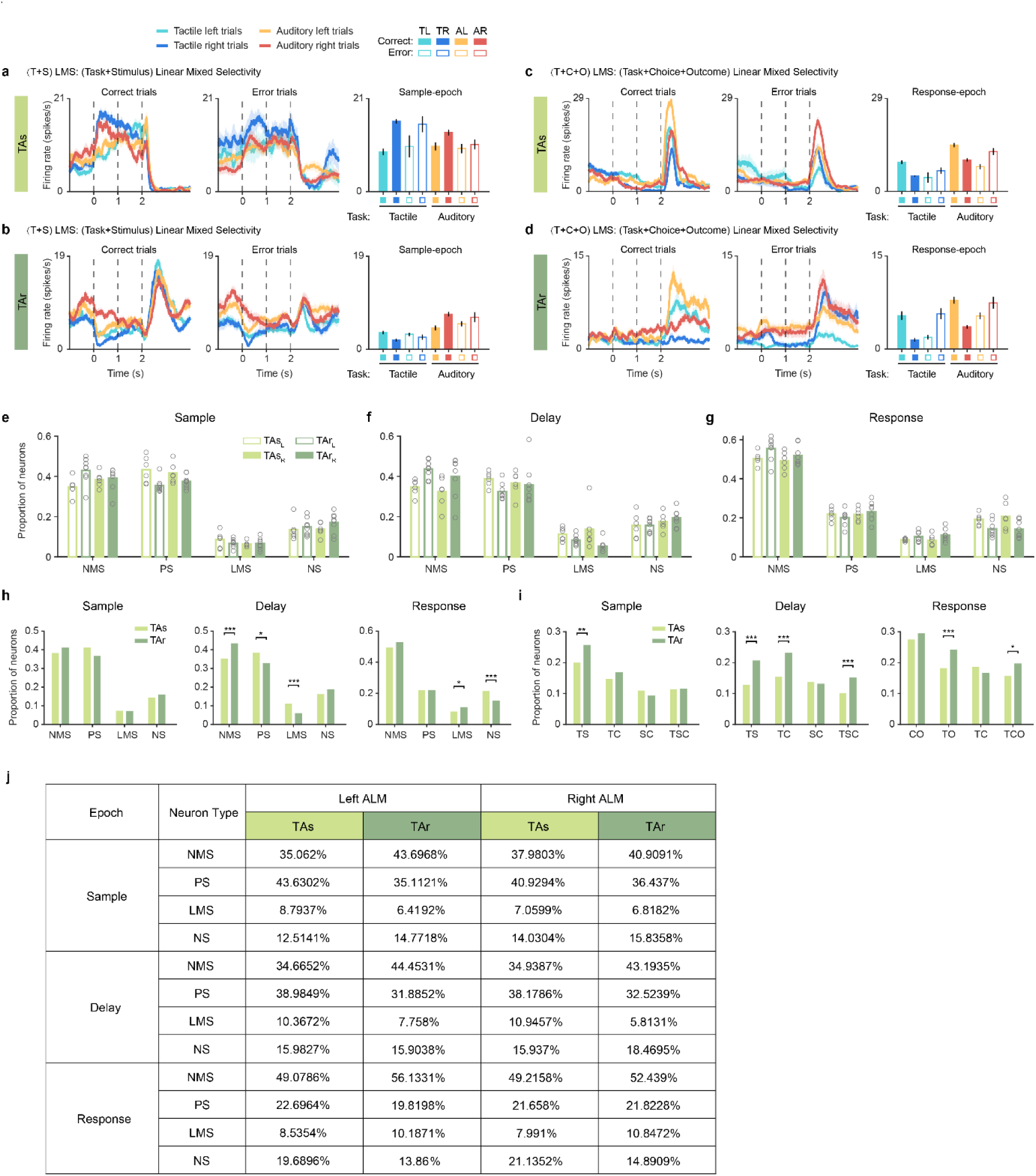
Extended analysis of neuronal mixed selectivity. **a**–**b**, Same as Fig. 4 a–b, but for example LMS neurons fitted sample-epoch firing rate with T, S and C. **c**–**d**, Same as Fig. 4 c–d, but for example LMS neurons fitted response-epoch firing rate with T, C and O. **e**, Proportions of NMS, PS, LMS and NS neurons across different epochs in the left and right ALM of individual mice. Open circle, individual animal; Bar, mean across animals. TAs, n = 6; TAr, n = 8. **f–g**, Same as Fig. 4e–f, but for the right ALM. \**P*<0.05, \*\**P*<0.01, \*\*\**P*<0.001, two-sided permutation test, Bonferroni correction for multiple comparison. **h**, Table showing the proportions of different neuron types in the left and right ALM of TAs and TAr mice.

**Extended Data Fig. 10.**
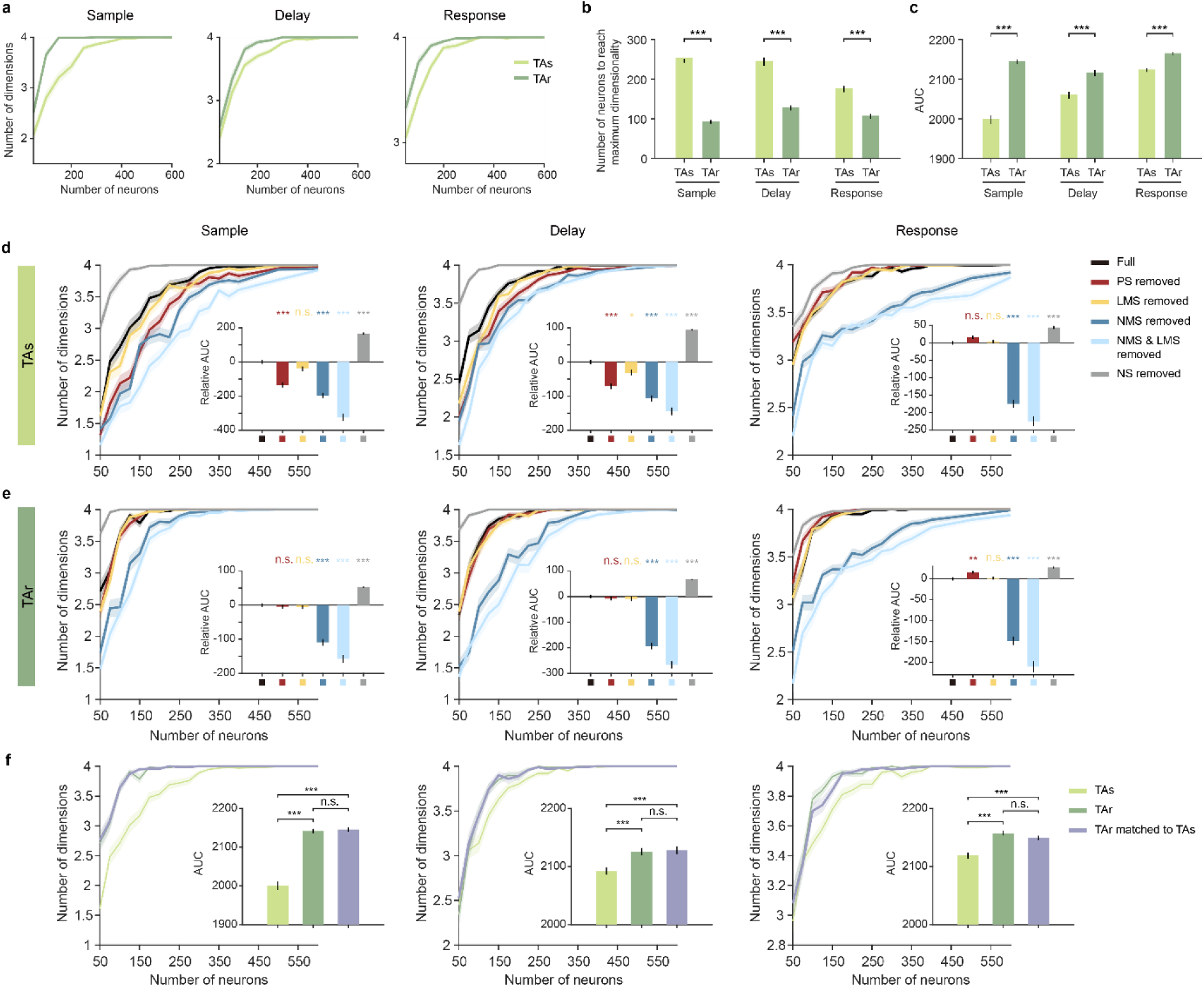
Representational dimensionality analysis of neurons from the right ALM. **a**– **c**, Same as Fig. 5 g–i, but for quantification of dimensionality of neurons in the right ALM. **d**–**e**, Same as Fig. 4 g–h but for the right ALM in TAs and TAr mice, respectively. Dimensionality of neural representations after removing different neuron types were analyzed. **f**, Same as Fig. 4 i but for the right ALM comparison. Matching proportions of neuron types in TAr to those in TAs barely influenced the number of dimensions (purple).

**Extended Data Fig. 11.**
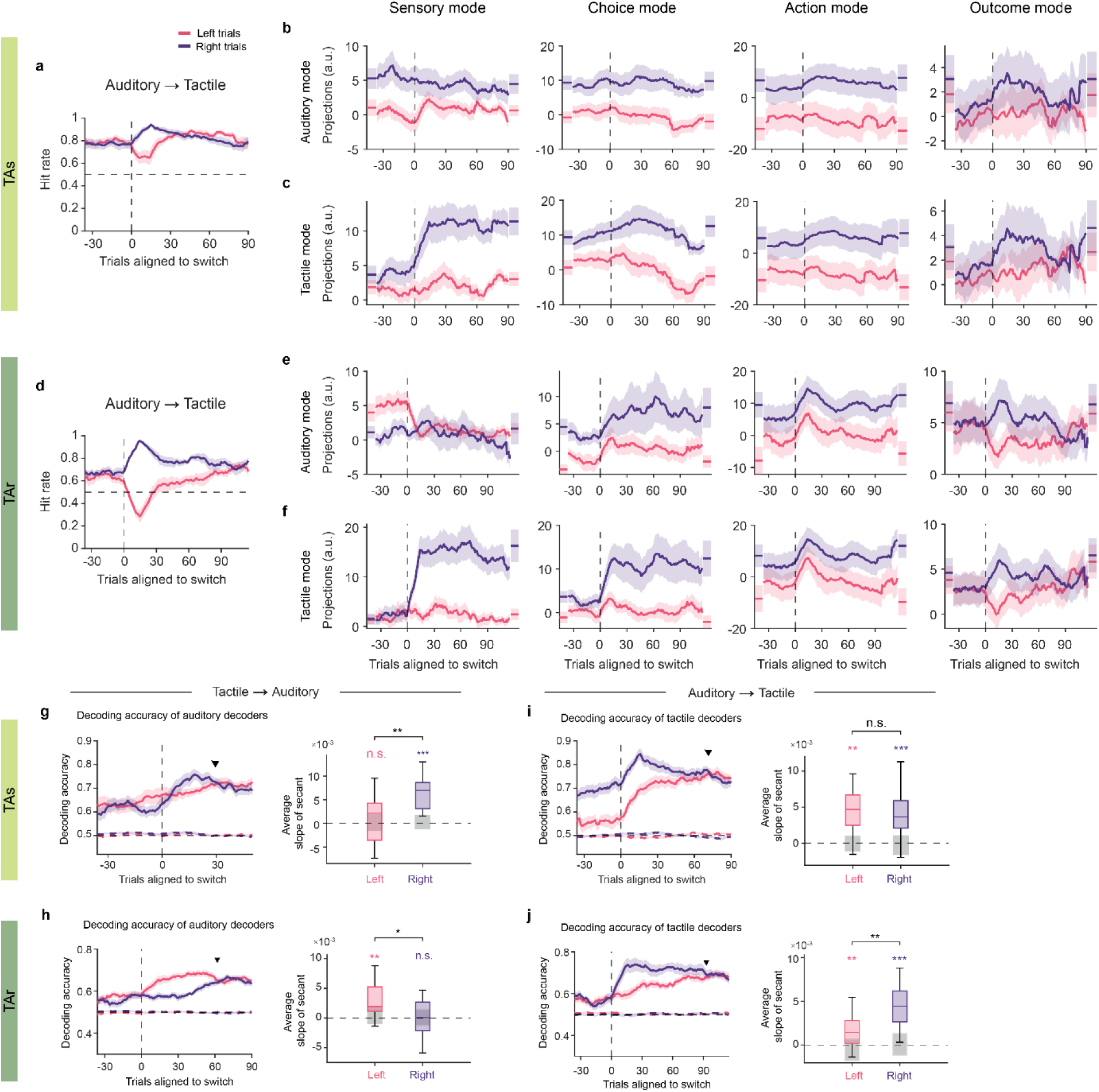
Extended analysis of neural reorganizations during task transition. **a**–**f**, Same as Fig. 6a–f but for auditory-to-tactile task transition. **a**, Behavioral performance of TAs mice transitioning from auditory to tactile task, mean ± s.e.m. across 18 sessions. **b**–**c**, Projections of single-trial activity along activity modes of auditory (pre-transition) task and tactile (post-transition) task, respectively. **d**–**f**, Same as a–c but for TAr mice, 19 sessions. Neural activity along the sensory mode changed immediately after transition in both contingencies, whereas projections along the choice and action modes exhibited remarkable differences between contingencies. **g**–**j**, Decoding choice using classifiers trained on activities in stable trials of the post-transition task during task transition. **g**, Decoding accuracy of auditory choice decoders in TAs mice during tactile-to-auditory transition. Left panel, decoding accuracy over trials aligned to transition onset, mean ± s.e.m. (bootstrap) across sessions. Solid line, decoder trained on true data; dashed line, null decoder trained on label-shuffled data. Downward triangle, recovery window. Right panel, quantification of the average change in decoding results from the switch to recovery window (Methods) Box plot shows the median and the 25th and 75th percentiles; whiskers represent 5th to 95th percentile range. Grey box, 5th to 95th percentile range of the null model. *P* value, two-sided Wilcoxon signed rank test; colored labels, comparison between true with null models; black labels, comparison between left and right trials of true data. \**P* < 0.05, \*\**P* < 0.01, \*\*\**P* < 0.001. **h**, Same as g but for TAr mice. **i–j**, Same as g–h but for auditory-to-tactile transition. Notably, choice decoding accuracy for post-switch trials with strong stimuli increased more rapidly after task switching.

**Extended Data Fig. 12.**
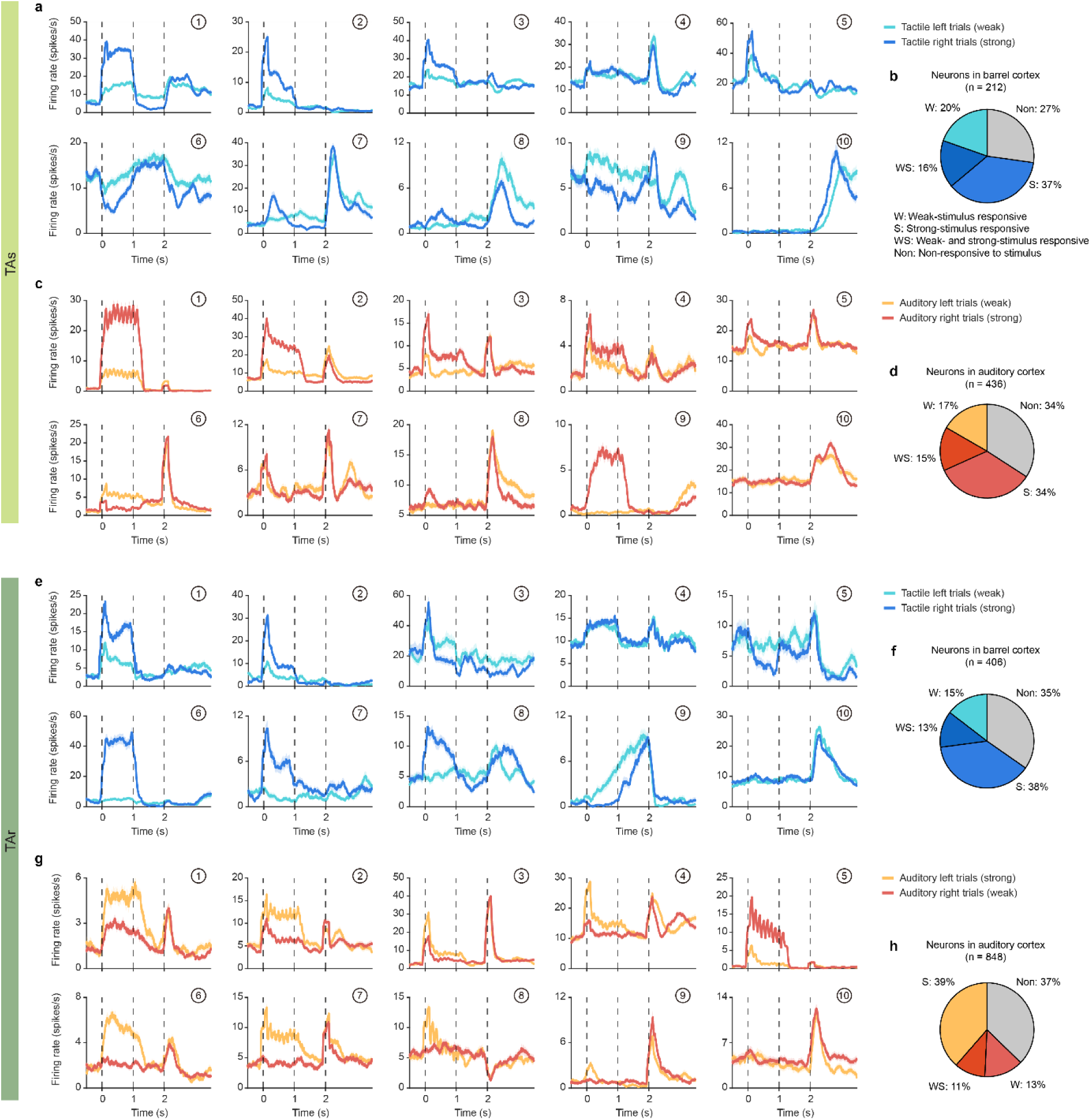
Neural activity in the primary somatosensory barrel cortex and auditory cortex. **a**, Example neurons in the barrel cortex of TAs mice. Neurons were classified by their early-sample responses (0–0.5 s from sample onset) relative to pre-sample baseline (−0.5–0 s) using a Mann–Whitney U test (*P* < 0.05) in tactile left trials with weak stimuli and tactile right trials with strong stimuli. Weak-stimulus responsive (W) neurons showed significant modulation for weak but not strong stimuli (example unit 9). Strong-stimulus responsive (S) neurons showed the opposite pattern, responding to strong but not weak stimuli (example units 7, 8). Neurons significantly modulated in both conditions were labeled weak-and strong-stimulus responsive (WS; example units 1–6). WS neurons typically fired more strongly to strong stimuli, though in some cases responses were comparable across strengths (unit 4). **b**, Proportions of stimulus-responsive and non-responsive neurons in the barrel cortex. **c**, Example neurons in the auditory cortex of TAs mice. Units 1–7 responded to both weak and strong stimuli; unit 8 and 9 responded only to the strong stimulus. **d**, Proportion of each type of neurons in the auditory cortex. **e**–**h**, Same as a–d but for TAr mice. Units 1–5 in e and g are responsive to both weak and strong stimuli, units 6–8 in e and units 6–9 in g are strong-stimulus responsive.

